# Conformational signatures of native ligand and pharmacochaperone binding in rhodopsin

**DOI:** 10.64898/2026.03.03.709294

**Authors:** Zaiddodine Pashandi, Joseph T. Ortega, Masaru Miyagi, Marcin Golczak, Beata Jastrzebska

## Abstract

Rhodopsin misfolding underlies rhodopsin-linked retinitis pigmentosa, and small-molecule pharmacochaperones represent a promising therapeutic strategy. However, the mechanisms by which these compounds interact with and stabilize rhodopsin remain poorly understood. Here, we combine backbone amide hydrogen-deuterium exchange mass spectrometry (amide HDX-MS), histidine-specific HDX (His-HDX), protein structure network (PSN) analysis, molecular docking, and functional spectroscopy to define ligand-induced conformational signatures in this receptor elicited by three non-retinoid small molecules, quercetin, myricetin, and the chromenone CR5, and to compare them with those of the native chromophore 11-*cis*-retinal. Binding of 11-*cis*-retinal to ligand-free opsin establishes a benchmark orthosteric conformational signature, characterized by strong backbone HDX protection across TM4-TM7 and adjacent loops, suppression of EX1-like hydrogen-deuterium exchange kinetics at the N-terminal ends of TM1 and TM4, and reorganization of PSN hubs that stabilizes an inactive-state residue interaction network. All three non-retinoid ligands generate HDX footprints that closely track this retinal-induced pattern within the chromophore pocket, consistent with direct orthosteric engagement, but they confer weaker and ligand-specific stabilization. Among them, quercetin most closely reproduces the retinal-like backbone protection and His-HDX microenvironment changes, whereas myricetin and CR5 only partially recapitulate retinal-induced stabilization and redistribute conformational flexibility toward TM1 and intradiscal regions, without fully suppressing EX1-like gating. In addition, all three compounds induce weak cytoplasmic allosteric effects in retinal-bound rhodopsin, indicating secondary interactions in addition to a primary orthosteric mechanism. Together, these results provide the first residue-level experimental framework for understanding the differential pharmacochaperoning capacity of non-retinoid ligands and highlight key conformational principles for future optimization of opsin stabilizers.

## INTRODUCTION

Rhodopsin, a G protein-coupled receptor exclusively expressed in rod photoreceptors, is the primary mediator of light detection that initiates electrochemical signaling to the brain ^1,2^. In its native state, this seven-transmembrane receptor contains a covalently bound 11-*cis*-retinal chromophore, which serves as the light sensor. Upon photon absorption, 11-*cis*-retinal isomerizes to its all-*trans* configuration inducing conformational changes in the protein scaffold that shift rhodopsin from its inactive to active state ^3,4^. Ultimately, all-*trans*-retinal dissociates from the binding pocket, leaving behind unliganded opsin. The released all-*trans*-retinal must then be converted back to the 11-*cis* configuration through the visual cycle to regenerate functional rhodopsin ^5^. More than 200 naturally occurring mutations in the rod opsin (*RHO*) gene have been identified, many of which cause retinitis pigmentosa (RP) and other degenerative retinal disorders that ultimately lead to vision loss ^6,7^. Currently, no approved treatment exists to prevent or slow the progression of *RHO*-related RP. Pharmacochaperone therapy has emerged as a promising strategy, as small molecules capable of stabilizing misfolded rod opsin mutants can enhance their proper folding, trafficking to the plasma membrane, and restore function ^8–10^. This therapeutic approach has gained significant interest and is being actively pursued by several research groups, including ours. Although retinal rescue of misfolded mutants has been demonstrated *in vitro* using the natural ligand 11-*cis*-retinal and its analogs, their chemical instability, and photoreactivity limit their therapeutic potential ^8,11,12^. Thus, non-retinoid small molecule pharmacochaperones represent a more promising alternative. Recently, we have identified several such non-retinoid pharmacochaperone molecules, including flavonoids such as quercetin and myricetin ^13–15^, CR5 chromenone ^16^, and synthetic JC3 and JC4 compounds ^17^ that bind to unliganded opsin with high affinity and enhance plasma membrane targeting of misfolding rod opsin mutants in cultured cells. These compounds also demonstrated therapeutic potential in mouse models of retinal degeneration ^14–17^.

Despite their therapeutic effects, the molecular and structural basis underlying the actions of these pharmacochaperones remains poorly understood. In the present study, we employed mass spectrometry-based approaches such hydrogen-deuterium exchange (HDX) in conjunction with mass spectrometry (MS) to analyze conformational rearrangements in opsin upon binding of flavonoid and chromenone compounds to the unliganded receptor. HDX is a powerful method for probing protein conformational dynamics in response to specific stimuli or environmental changes. Traditional amide-HDX studies have provided valuable insights into solvent accessibility and ligand-induced conformational changes in the membrane proteins, including rhodopsin ^4,18,19^.

It enables the assessment of backbone-wide protein structural dynamics but is inherently insensitive to side-chain chemistry and lacks residue-level spatial resolution. In contrast, histidine (His)-HDX as established previously ^20^, provides site-specific information on local electrostatic environments and solvent accessibility. However, its structural coverage is restricted to regions containing His residues. Thus, the complementary application of amide-HDX and His-HDX provides a comprehensive view of global backbone dynamics while simultaneously offering site-specific insights into local structural features surrounding His residues. Both amide-HDX and His-HDX revealed changes in regions adjacent to the binding site upon flavonoids and CR5 engagement with ligand-free rod opsin, consistent with orthosteric binding. However, motivated by prior reports of allosteric modulation of rhodopsin by flavonoids ^21,22^ and other compounds ^23^, we also examined ligand-induced structural rearrangements in retinal-bound rhodopsin and observed weak allosteric modulation of its conformation by these compounds. Our findings are consistent with UV-Vis acid denaturation measurements indicating low affinity binding with dissociation constants (K_d_) in the micromolar range.

Nevertheless, our findings support orthosteric binding of non-retinoid pharmacochaperones as a primary strategy for correcting rod opsin misfolding, while revealing weaker allosteric modulation whose physiological significance requires further investigation.

## RESULTS

### Orthosteric Ligand-Induced Structural Rearrangements in Rod Opsin - Insights from Amide-HDX

Amide-HDX is a convolution of solvent accessibility, protein backbone dynamics, and intrinsic amide H/D exchange rates ^24^. However, it needs a consistent regime of protein digestion and peptide separation which is still challenging in the case of membrane proteins, containing multiple posttranslational modifications. By systematically varying pepsin concentration and digestion time, we have established robust peptide maps for affinity-purified unliganded rod opsin and 11-*cis*-retinal-bound rhodopsin. Approximately 70-80% sequence coverage was achieved with 10 min pepsin digestion, which was selected as the optimal condition for subsequent HDX experiments following ligand treatment (**Fig. 1**). Under these experimental conditions, individual peptides exhibited deuterium uptake values ranging from 0 to 30% of maximal hypothetical deuterium uptake (**Δ**% deuterium uptakes are summarized in **Supplementary Table 1**). Both the sequence coverage and uptake range were comparable to previously reported HDX peptide maps for rod opsin (**Fig. 1**) ^4,25,26^.

**Figure 1.**
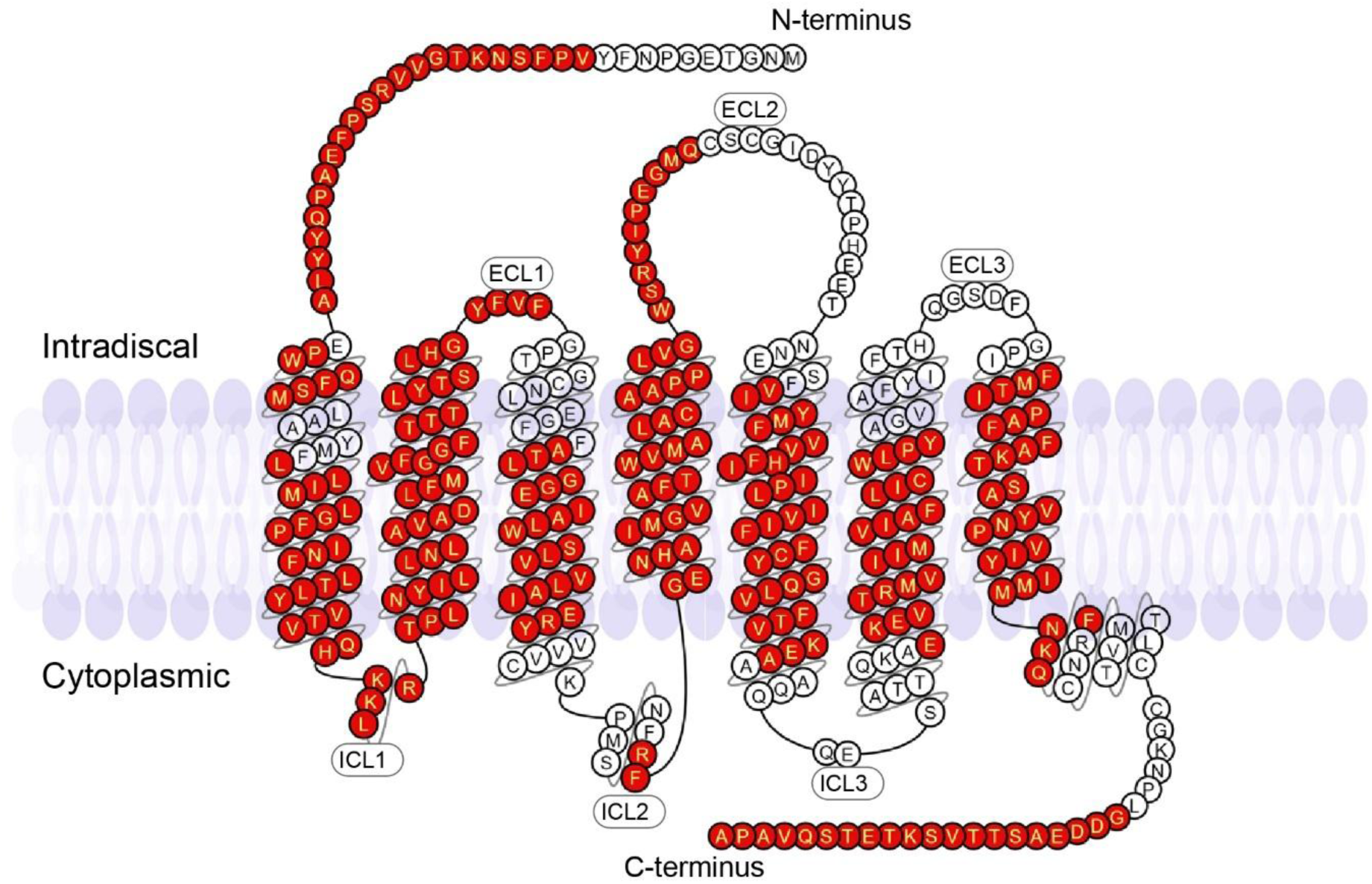
Peptide coverage of the rod opsin upon pepsin digest determined by mass spectrometry. Shown is a two-dimensional map of rod opsin in which amino acid residues covered by identified peptides in either H₂O or D₂O conditions are highlighted in red.

Overall, ligand-free opsin exhibited increased deuterium uptake compared with dark-state rhodopsin, indicating greater conformational flexibility in the absence of the endogenous ligand. HDX difference mapping between opsin and rhodopsin revealed substantial conformational changes upon binding of 11-*cis*-retinal within the orthosteric pocket (**Fig. 2A**). Specifically, transmembrane helices surrounding the retinal-binding site became less solvent accessible following 11-*cis*-retinal binding, consistent with ligand-induced stabilization and structural rearrangement ^4^. The largest decreases in deuterium uptake (>10%) were observed for peptide fragments corresponding to transmembrane helices (TM) 4, 5, 6, and 7, particularly in regions proximal to the β-ionone ring location (**Supplementary Table 1**).

**Figure 2.**
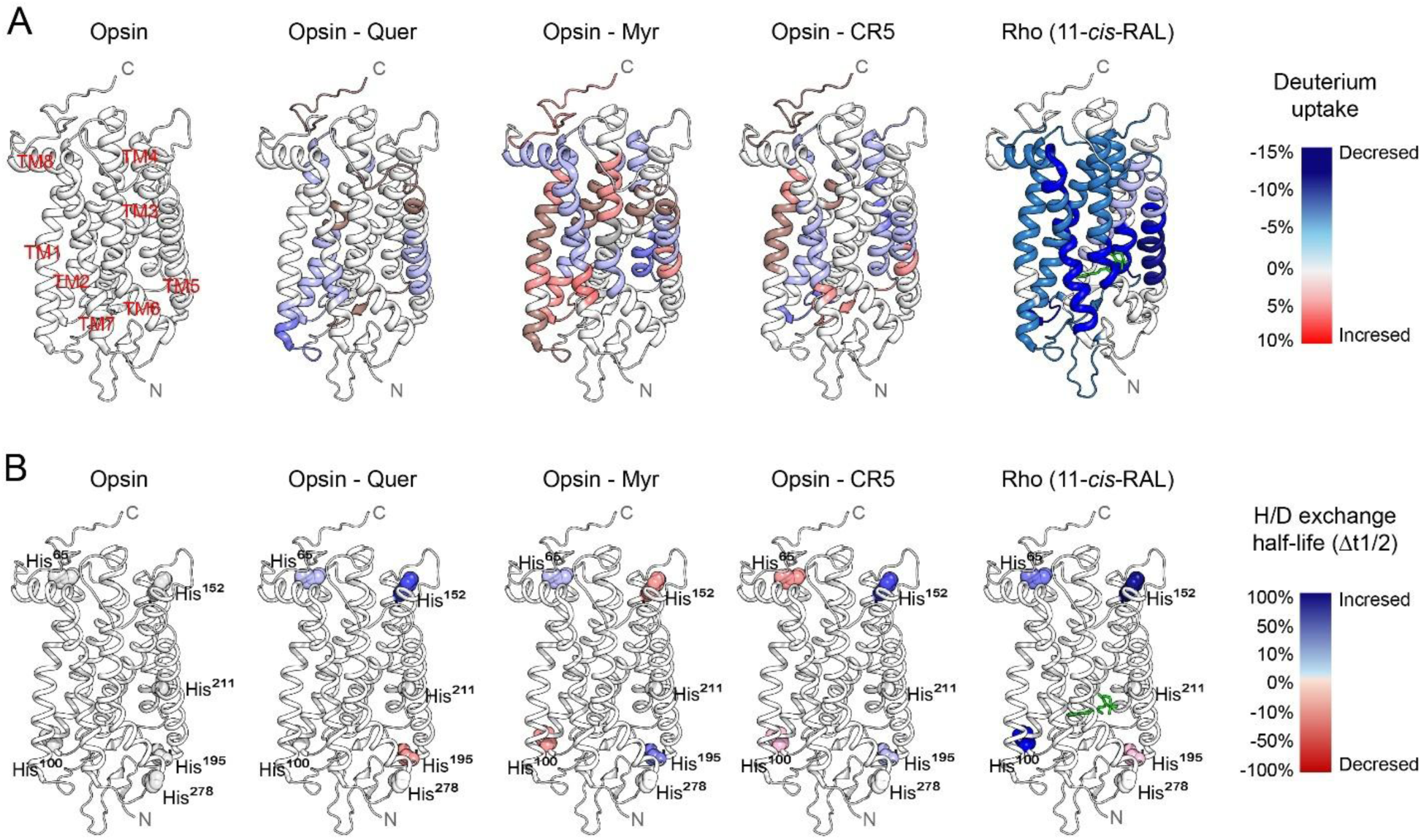
Orthosteric conformational signatures of rod opsin-ligand complexes. **A**, Amide-HDX difference map of rod opsin-ligand complexes. The differential percent deuteration for ligand-free opsin upon binding of each ligand. The percent deuteration of the compound-treated opsin was subtracted from the percent deuteration of the untreated opsin and displayed on the surface of the receptor as a heat map. The blue color displays the percentage of deuterium uptake reduction (decreased solvent accessibility). The red color shows the percentage increase in deuterium uptake (increased solvent accessibility). Transmembrane helices (TM) are indicated by red labels in the opsin structure. 11-*cis*-retinal is shown with green sticks in the rhodopsin (Rho) structure. **B**, His-HDX difference map of rod opsin-ligand complexes. The differential percent half-life for ligand-free opsin after incubation with each ligand. The deuteration half-life of the opsin-ligand complexes was subtracted from the deuteration half-life of the untreated opsin and displayed on the corresponding histidine residues as a heat map. The blue color displays differential increase in His-HDX half-life (decreased solvent accessibility). The red color shows differential decrease in His-HDX half-life (increased solvent accessibility).

Incubation of apo-opsin with non-retinoid ligands, including quercetin, myricetin, and chromenone CR5 produced HDX patterns that largely recapitulated those observed upon 11-*cis*-retinal binding, despite ligand-specific differences in the magnitude of deuterium uptake. This similarity in the HDX difference maps suggests that these flavonoids and CR5 bind within the orthosteric binding site (**Fig. 2A**). For all three ligands, decreased deuterium uptake was observed across multiple transmembrane helices such as TM2 and TM4-TM7, with the most pronounced effects localized to peptides surrounding the orthosteric binding site. However, in contrast to 11-*cis*-retinal, the overall extent of ligand-induced stabilization was less pronounced and largely confined to regions adjacent to the orthosteric cavity. Notably, treatment with chromenone CR5, and more prominently with myricetin, also resulted in increased deuterium uptake in TM1 and other regions distal to the orthosteric pocket, indicating ligand-specific conformational effects beyond the retinal-binding site.

### Orthosteric Ligand-Induced Changes in Histidine Microenvironment Detected via His-HDX

To gain site-specific insight into local electrostatic environments and solvent accessibility, His-HDX analysis was performed on opsin incubated with flavonoids or CR5, and the results were compared with 11-*cis*-retinal-bound rhodopsin. Six His residues distributed throughout the opsin structure served as intrinsic probes to detect structural differences among ligand-free, retinal-bound, and non-retinoid ligand-bound states. The half-life (*t_1/2_*) of each His residue reflected changes in its local microenvironment associated with distinct receptor conformations. Six unique peptides, each containing a single His residue, are listed in **Supplementary Table 2**. Analysis of His-HDX kinetics revealed clear differences in *t*_1/2_ for His^65^, His^100^, His^152^, and His^195^ when comparing opsin to 11-*cis*-retinal-bound rhodopsin and its complexes with flavonoids and CR5, while the *t*_1/2_ value for His^211^ and His^278^ could not be reliably determined under these experimental conditions. Binding the natural ligand to the orthosteric site increased *t*_1/2_ values of His^65^, His^100^, and His^152^, consistent with structural stabilization of these regions, accompanied by a modest decrease in *t*_1/2_ of His^195^ (**Fig. 2B**). Non-retinoid ligands produced both retinal-like and ligand-specific His-HDX signatures. Upon quercetin binding to unliganded opsin, the changes in the *t*_1/2_ for His^65^, His^152^, and His^195^ followed the same trends observed with 11-*cis*-retinal binding, whereas no significant change was detected for His^100^. In contrast, myricetin binding produced a retinal-like effect at His^65^, while His^100^, His^152^, and His^195^ exhibited changes in the opposite direction. For CR5-bound opsin, the change in the *t*_1/2_ at His^152^ was consistent with the effect of 11-*cis*-retinal, while His^65^, His^100^, and His^195^ displayed opposite trends. Together with the amide-HDX data, these results show that non-retinoid ligands partially recapitulate retinal-induced conformational stabilization while introducing distinct, ligand-specific modes of opsin modulation.

### Protein Structure Network (PSN) analysis

To understand how ligand binding reorganizes the global network of interactions within opsin, we performed PSN analysis, which provides a detailed map of inter-residue interaction networks ^27,28^. Since the structure of opsin bound to non-retinoid ligands is not determined experimentally, here we compare the structure network of ligand-free opsin (PDB ID: 3CAP) versus 11-*cis*-retinal bound rhodopsin (PDB ID: 1U19), and a previously generated MD simulation ensemble of opsin-quercetin complex ^15^. Notably, the PSN analysis showed strong agreement with the HDX pattern observed upon 11-*cis*-retinal binding (**Fig. 3A-B**) and quercetin binding (**Supplementary Fig. 1**). The amino acids and peptide segments contributing to MetaPath changes following 11-*cis*-retinal binding (**Fig. 3B**, orange links), together with their corresponding HDX signatures highlighted in **Fig. 3A**.

**Figure 3.**
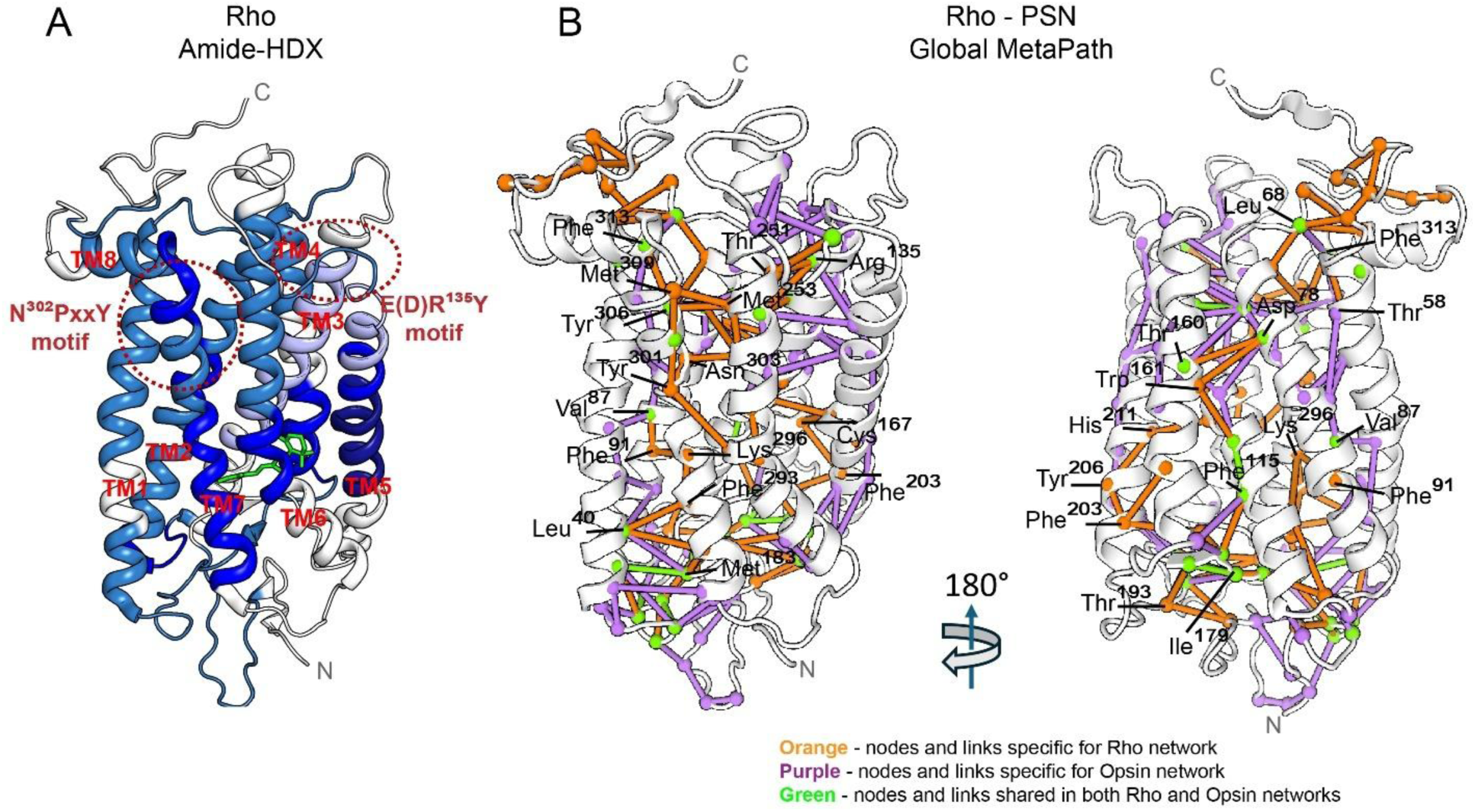
Structural amide-HDX pattern for 11-*cis*-retinal-bound rhodopsin and corresponding Protein Structure Network (PSN) models. **A**, The structural rearrangement model for rhodopsin (Rho) amide-HDX is shown in left panel. The intensity of blue color displays the percentage of deuterium uptake reduction (decreased solvent accessibility). Location of NPxxY, and E/DRY motifs are indicated by the red dashed ovals. 11-*cis*-retinal is shown with green sticks in the Rho structure. **B**, The Global MetaPath derived from PSN analysis. Nodes and links specific to ligand-free opsin (PDB ID: 3CAP) and 11-*cis*-retinal-bound Rho (PDB ID:1U19) networks are represented by purple and orange color respectively. Nodes and links shared by both networks are colored green. Generation of new MetaPaths after 11-*cis*-retinal binding (orange color) are in good concordance with change in percentage of deuterium uptake across transmembrane region, suggesting corresponding network and structural rearrangement following endogenous ligand binding.

These amino acids that represent formation of unique links include Glu^113^, Glu^122^, Leu^125^, and Ser^127^ from TM3; Trp^161^ and Cys^167^ from TM4; Phe^203^, Tyr^206^, and His^211^ from TM5; Met^253^, Met^257^, Phe^261^, Cys^264^, and Trp^265^ from TM6; Thr^289^, Phe^293^, Lys^296^, Tyr^301^, Asn^302^, Ile^305^, and Met^309^ from TM7; and Phe^313^ and Cys^316^ from TM8; as well as His^65^ and Lys^67^ from intracellular loop 1 (ICL1); Phe^103^ from extra cellular loop 1 (ECL1); and Glu^181^, Ser^186^, Ile^189^, and Thr^193^ from ECL2. Similarly, PSN comparisons between unliganded and quercetin-bound opsin (**Fig. S1**) revealed putative amino acids and quercetin-mediated interaction networks that likely underline the observed reductions in backbone HDX across TM1 (Ala^40^ and Tyr^43^), TM2 (Asn^73^, Tyr^74^, Ala^80^, Phe^88^, Phe^91^, Tyr^96^, Thr^97^, and Ser^98^), TM5 (Tyr^206^, Val^210^, and His^211^), TM7 (Ile^290^, Phe^293^, Phe^294^, Lys^296^, Thr^297^,Ser^298^, Tyr^301^, and Asn^302^), and TM8 (Asn^310^ and Met^317^); as well as ICL1 (Thr^70^ and Pro^71^). Together, these analyses illustrate how local rearrangements within the binding pocket propagate to neighboring segments and more distal regions, including the NPxxY and E/DRY motifs, revealing how pharmacochaperones globally rewire receptor interaction networks to stabilize the receptor. This framework provides a mechanistic explanation for why chemically distinct ligands produce divergent long-range conformational signatures.

### Signature of Rhodopsin Allosteric Modulation

Previous studies suggest that rhodopsin can be modulated allosterically ^23,29^, and that flavonoids may interact with this receptor at allosteric sites located in both cytoplasmic and extracellular regions ^21,22,30^. However, these interactions are typically weak, with binding affinities in the micromolar range ^23,29^. To explore the structural signatures of these weak allosteric interactions upon binding of non-retinal ligands tested in this study, first we applied UV-Vis spectroscopy and acid denaturation assay. This assay allows to monitor acid-induced disruptions of rhodopsin conformation, which induces a decrease of the absorption maximum at 500 nm and an appearance of the absorption peak at 440 nm, reflecting destabilization of the protein tertiary structure and Schiff base deprotonation (**Fig. 4A**) ^21,31,32^. At a fixed concentration of ligand and acid, all three compounds accelerated the decay of the 500 nm absorbance peak, indicating enhanced acid-induced destabilization of dark-state rhodopsin and supporting allosteric modulation by these non-retinoid ligands (**Fig. 4B**). Moreover, concentration-dependent measurements revealed their weak micromolar binding affinities (K_d_ = 1.35±0.25, 1.60±0.19, and 3.57±0.54 µM), affinities ranked as myricetin > CR5 > quercetin, consistent with an allosteric mode of action (**Fig. 4C**).

**Figure 4.**
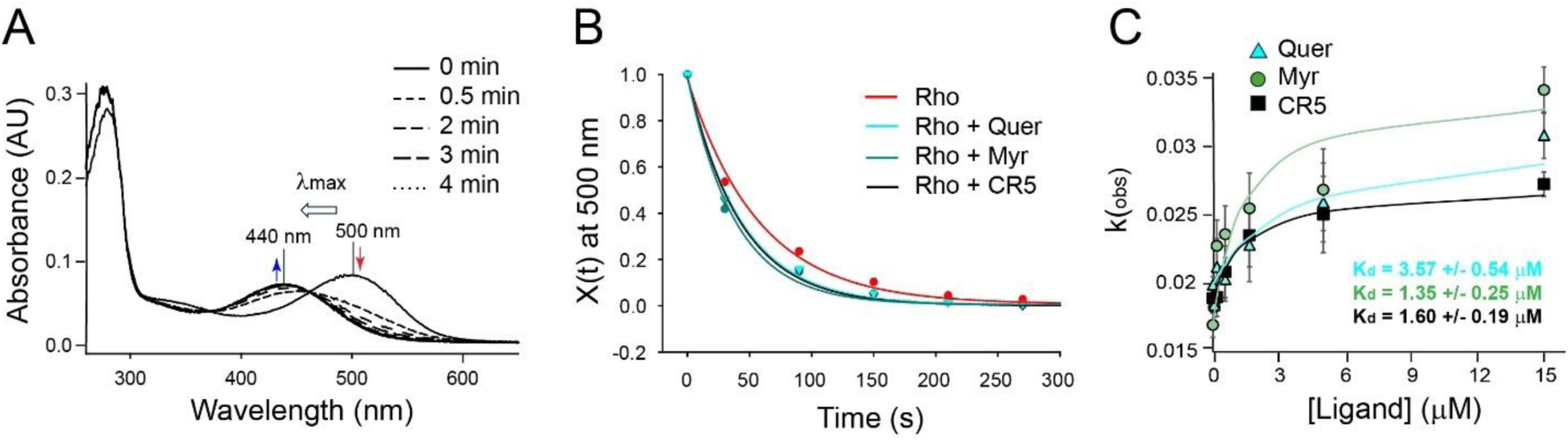
Effect of allosteric interaction on the rate of acid denaturation. **A**, The representative changes in absorbance at 500 nm and 440 nm for rhodopsin (Rho) following acid treatment (pH < 3). **B**, Time-dependent decay of rhodopsin absorbance at 500 nm measured in the absence and presence of a non-retinoid ligand at a concentration equal to its K_d_, under 30 mM acid conditions. **C**, Observable rate constant for native fraction decay (k_obs_) plotted as a function of ligand concentration (0, 0.062, 0.19, 0.56, 1.67, 5, and 15 µM) for myricetin (green circles), quercetin (cyan triangles), and CR5 (black squares). The values of dissociation constants (K_d_) for the allosteric interaction, calculated using Equation (III) (Experimental Methods), are indicated in the figure.

To investigate conformational rearrangements in rhodopsin upon binding of these non-retinoid ligands we applied HDX-MS analyses. Consistent with our UV-Vis spectroscopy results, we found that, even when the orthosteric binding site is occupied by the native ligand 11-*cis*-retinal, each non-retinal compound induces detectable structural rearrangements in the receptor (**Fig. 5A**), supporting their role in allosteric modulation. Among the ligands tested, myricetin and CR5 produced the most pronounced effects, increasing deuterium uptake across multiple transmembrane helices, indicative of enhanced conformational dynamics. Decreases in deuterium uptake were more localized, primarily at residues 85-90 and the cytoplasmic end of TM4. In contrast, quercetin elicited more modest, spatially restricted changes, including slight decreases near the cytoplasmic end of TM7 and a segment of TM5 (residues 209-214), along with a minor increase near the extracellular end of TM1. Consistent with the amide-HDX results, incubation of 11-*cis*-retinal-bound rhodopsin with each compound revealed additional structural changes around histidine side chains, further supporting their potential allosteric effects (**Fig. 5B**). Quercetin binding led to a decrease in the His-HDX *t*_1/2_ for His^65^, His^100^, and His^152^, whereas myricetin and CR5 increased *t*_1/2_ values at His^65^ and His^195^, respectively, accompanied by decreases at the remaining His residues. These ligand-specific patterns highlight distinct modes of allosteric modulation, collectively supporting the conclusion that flavonoids and CR5 can differentially modulate the conformational dynamics of dark-adapted rhodopsin.

**Figure 5.**
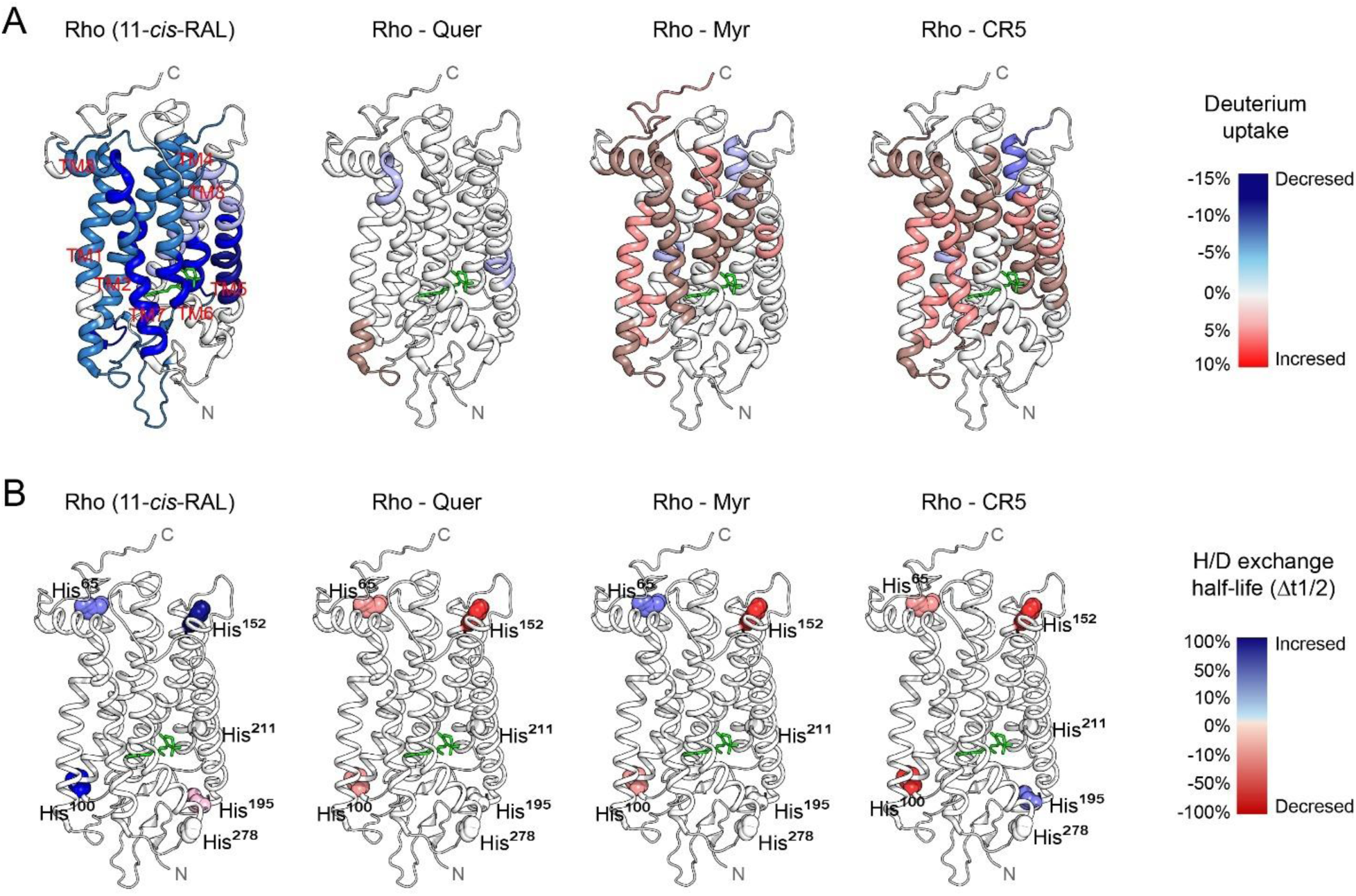
Allosteric conformational signatures of rhodopsin-ligand complexes. **A**, Amide-HDX difference map of rhodopsin-ligand complexes. Differential percent deuteration of rhodopsin (Rho), with the orthosteric pocket occupied by 11-*cis*-retinal, upon incubation with non-retinoid compounds. The percent deuteration of the compound-treated rhodopsin was subtracted from the percent deuteration of the untreated rhodopsin and displayed on the surface of the receptor as a heat map. The blue color displays the percentage of deuterium uptake reduction (decreased solvent accessibility). The red color shows the percentage increase in deuterium uptake (increased solvent accessibility). **B**, His-HDX difference map of rhodopsin-non retinoid ligand complexes. The differential percentage half-life calculated for rhodopsin upon incubation with non-retinoid compounds. The location of all 6 His residues is indicated to facilitate the orientation of the receptor’s 3D. The deuteration half-life of the rhodopsin-non-retinoid ligand complexes was subtracted from the deuteration half-life of the untreated rhodopsin and displayed on the corresponding His residues as a heat map. The blue color displays differential increase in His-HDX half-life (decreased solvent accessibility). The red color shows differential decrease in His-HDX half-life (increased solvent accessibility).

### Elucidation of Putative Amino Acid Residues Involved in Ligand Binding in Opsin and Rhodopsin

To further characterize the molecular interactions between quercetin, myricetin and CR5, and both orthosteric and allosteric sites, we performed molecular docking simulations of these compounds to opsin and rhodopsin using AutoDock Vina in the PyRx ^33–35^. Docking scores and interaction types for all ligand-receptor complexes are summarized in **Supplementary Table 3**, while the close-up views of the interaction networks for the best poses at the orthosteric and allosteric sites are shown in **Figs. 6** and **7**, respectively. As the coordination of Trp^265^ and Tyr^268^ differs between the inactive-like (PDB ID: 2I36) and active-like (PDB ID: 3CAP) conformations of apo-opsin, molecular docking was performed using both structures (**Fig. 6A-B**, respectively).

**Figure 6.**
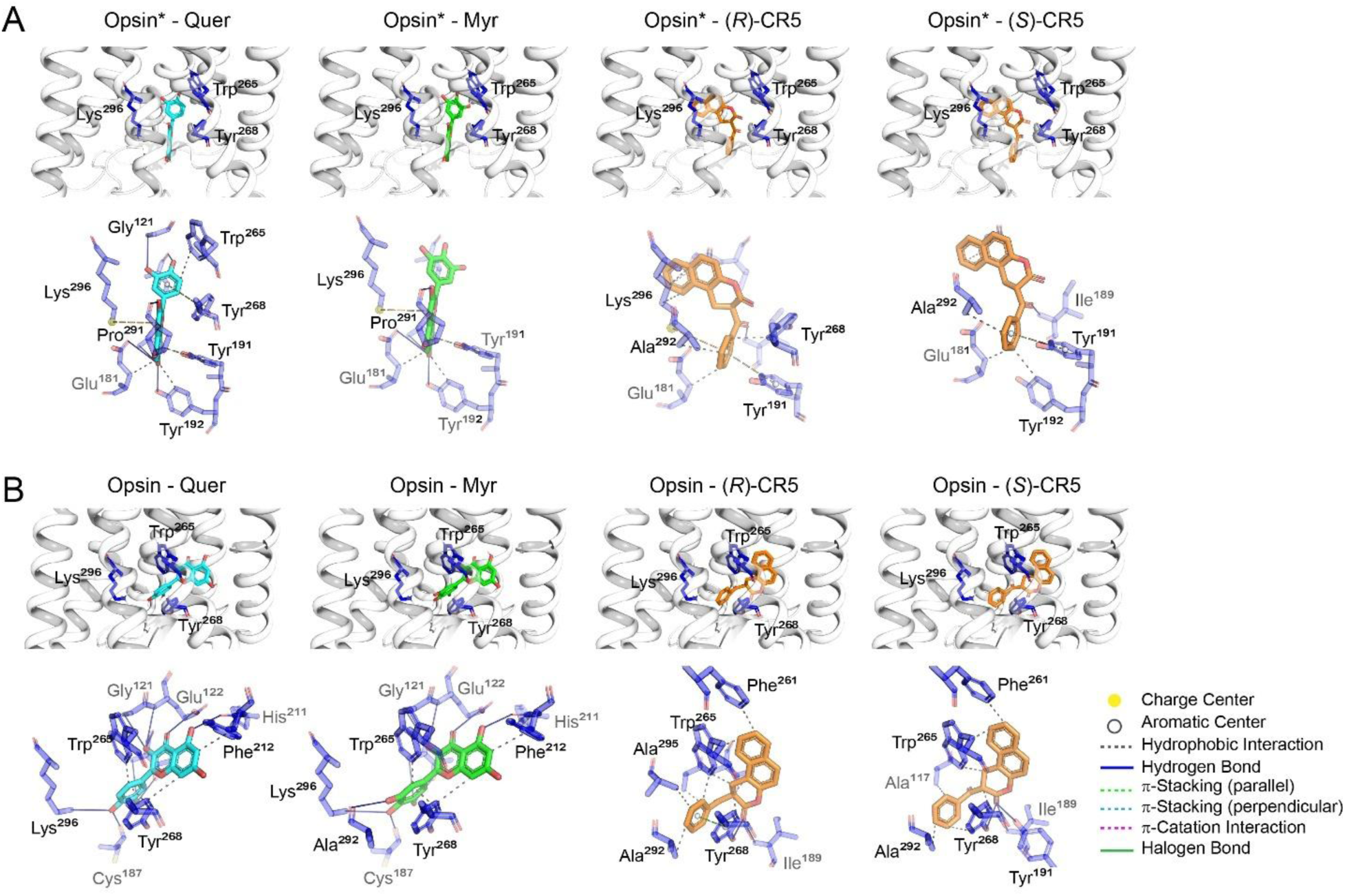
Local interactions between bovine rod opsin and flavonoids or CR5 within the orthosteric pocket. **A-B**, Molecular docking analysis of ligand interactions within the orthosteric pocket of rod opsin using structures of apo-opsin with active conformation (opsin*) (PDB ID: 3CAP) (**A**) and inactive conformation (opsin) (PDB ID: 2I36) (**B**). Docking was performed using AutoDock Vina in PyRx environment. The top panel shows the ‘zoomed-in’ view for the best docking poses for each ligand according to the docking scores. And the bottom panel depicts the predicted interactions between the ligand and amino acid residues within orthosteric pocket, identified with the Protein-Ligand Interaction Profiler (PLIP) web server.

**Figure 7.**
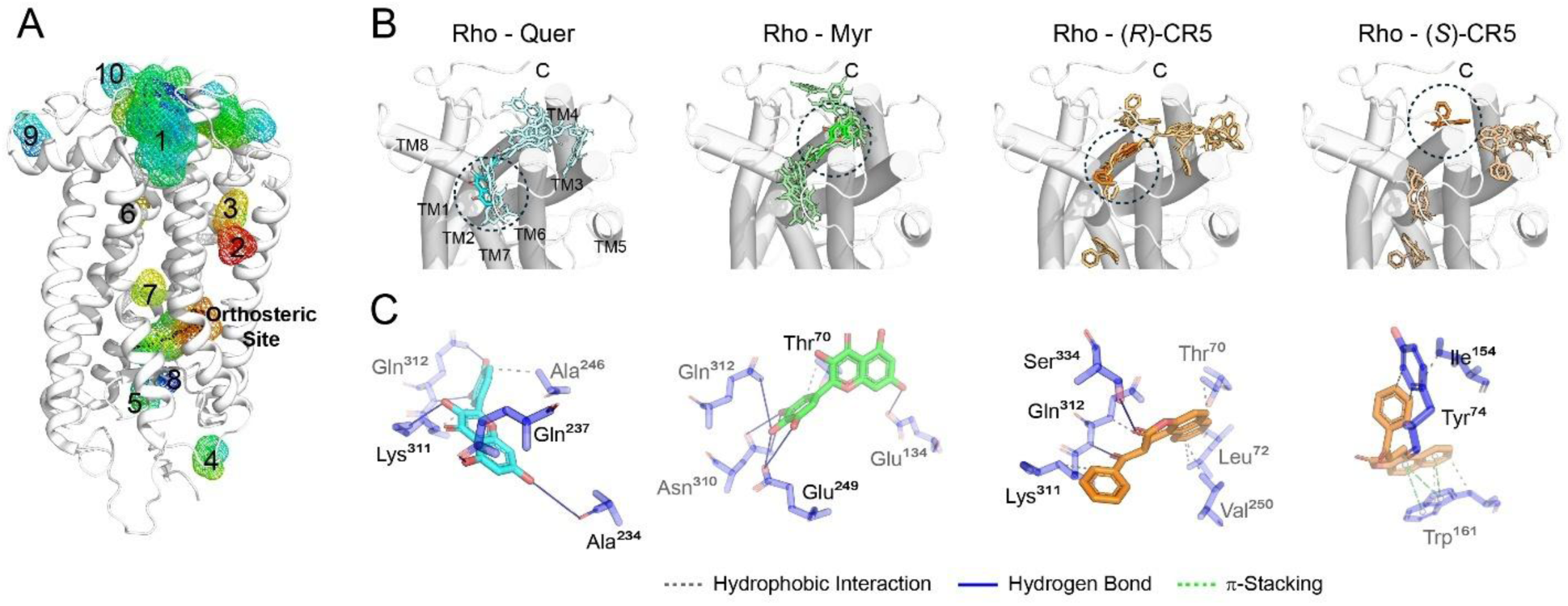
Allosteric interactions of flavonoids and CR5 with the cytoplasmic cavity of bovine rhodopsin. **A**, The allosteric cavities in rhodopsin predicted using CavitOmix plugin in PyMol with BioNeMo server for structure prediction. Contour colors are displayed based on hydrophobicity (lowest hydrophobicity: blue, to highest hydrophobicity: red). The top 10 predicted allosteric cavities are shown. The corresponding cavity volumes are as follows: 1 - 1880 Å^3^; 2 - 80 Å^3^; 3 - 68 Å^3^; 4 - 63 Å^3^; 5 - 53 Å^3^; 6 - 47 Å^3^; 7 - 43 Å^3^; 8 - 41 Å^3^; 9 - 49 Å^3^; 10 - 40 Å^3^. **B-C**, Molecular docking analysis of non-retinoid ligand interactions in the cytoplasmic cavity of dark state rhodopsin (Rho), (PDB ID: 1U19). Docking was performed using AutoDock Vina in PyRx environment. **B**, The ‘zoomed-in’ view of all docking poses identified for each ligand in the cytoplasmic region with the best-ranked poses highlighted by black dotted circles. **C**, Predicted interactions between the non-retinoid ligands and the amino acid residues in the cytoplasmic region, identified using the Protein-Ligand Interaction Profiler (PLIP) web server.

In the 3CAP model, all suggested poses for the ligands were located near Lys^296^ and are stabilized mostly by residues from TM3, TM6, TM7, and ECL2 (**Supplementary Fig. 2**), whereas in the 2I36 model, orthosteric poses exhibited variable orientations, extending into the β-ionone cavity to the vicinity of TM5 (**Supplementary Fig. 3**). A few poses for quercetin, myricetin, and CR5 were located outside the orthosteric pocket, but these showed lower predicted binding affinities (**Supplementary Fig. 3**). Using each initial model, the top-ranked docking poses of quercetin and myricetin fully overlapped and stabilized the receptor through nearly identical interactions. In the 3CAP model, the catechol phenyl ring of both flavonoid compounds was oriented toward Trp^265^ and Gly^121^, while the flavone core formed hydrogen bonds with Glu^181^, Tyr^191^, Tyr^192^, and Pro^291^, and engaged in a π-cation interaction with Lys^296^ (**Fig. 6A**). In contrast, in the 2I36 model, the flavonoids adopted a rotated orientation in which the catechol phenyl ring faced Lys^296^ and formed hydrogen bonds, whereas the flavone core occupied the β-ionone cavity and interacted with Glu^122^, His^211^ and Phe^212^ (**Fig. 6B**). For CR5, the *(R*)- and (*S*)-enantiomers showed substantial overlap within each model. However, a pronounced positional shift of the CR5 core was observed between the 3CAP and 2I36 structures. In the 3CAP model, the CR5 core formed hydrophobic interactions with Ala^117^, whereas in the 2I36 model it rotated toward the β-ionone cavity and interacted with Phe^261^ and Trp^265^ (**Fig. 6A-B**). Notably, the phenyl substituent of CR5 remained in a similar position in both models, consistently interacting with Ile^189^, Tyr^268^, and Ala^292^.

Since our HDX-MS analysis (**Fig. 5**) revealed that in addition to orthosteric binding, these ligands induce weak allosteric modulation of rhodopsin we performed computational prediction analysis using CavitOmix plugin in PyMol with BioNeMo mode and identified several potential allosteric cavities on the rhodopsin surface with the largest (volume 1882 Å^3^) located at the cytoplasmic site (**Fig. 7A, cavity 1**). To identify specific interactions between receptor and investigated non-retinoid ligands in this region, quercetin, myricetin, and CR5 were docked into this cytoplasmic cavity of dark-state bovine rhodopsin structure (PDB ID: 1U19). Quercetin and myricetin primarily occupied the central cavity formed between the cytoplasmic ends of the transmembrane helices TM2-TM4, TM6 and TM7, whereas both CR5 (*R*)- and (*S*)-enantiomers exhibited more dispersed binding distributions. Some CR5 poses indicate binding within the central cytoplasmic cavity, others along inter-helical grooves near the cytoplasmic ends (**Fig. 7B-C)**. The best-ranked pose of quercetin was stabilized via hydrogen bonding with Ala^234^ and Gln^237^ from ICL3, as well as Ly^311^ and Gln^312^ from TM7. The top-ranked pose of myricetin was located at the interface of TM2 (Thr^70^), TM3 (Glu^134^), TM6 (Glu^249^), and TM7 (Asn^310^ and Gln^312^), where it formed multiple hydrogen-bonding and hydrophobic interactions. In both cases the best-scoring poses align well with regions where the HDX difference maps show protection (**Fig. 5A**). For (*R*)-CR5, the best docking pose resided in a cleft between the cytoplasmic ends of TM2 and TM6 and TM7, stabilized by hydrophobic interaction with Thr^70^, Lue^72^, and Val^250^, and hydrogen bonds with Gln^312^ and Ser^334^. In contrast top-ranked pose of (*S*)-CR5 preferentially occupied a groove between TM2 and TM4 near their cytoplasmic ends engaging in π-π stacking with Trp^161^ and hydrophobic interactions with Tyr^74^ and Ile^154^. Although the top scoring docking poses of both (*R*)- and (*S*)-CR5 are located somewhat in distal to the site predicted by HDX, several lower-ranked pose clusters were found near the HDX-defined region of TM4, supporting the idea that CR5 samples a small ensemble of closely related cytoplasmic binding modes.

### EX1-Like HDX at the Flexible Ends of TM1 and TM4 Reveals Rare Cooperative Opening Events

In HDX-MS, EX1 kinetics typically arise when a protein segment undergoes relatively rare but cooperative opening events, corresponding to local or global dynamics that simultaneously expose all exchangeable backbone amides, followed by rapid deuterium incorporation before the structure recloses. Under these conditions, the observed exchange behavior is governed by the opening rate constant (k_open_), resulting in bimodal isotopic envelopes that represent distinct closed (low exchange) and open (high exchange) receptor sub-populations ^36,37^. In contrast, EX2 kinetics reflect frequent opening and closing fluctuations in proteins, with deuterium incorporation limited by the intrinsic chemical exchange rate, producing a single isotopic envelope that shifts gradually over time ^38,39^.

Among all peptide segments analyzed by backbone amide-HDX, two regions, EPWQFSML (residues 33-40) at the N-terminal end of TM1 (**Fig. 8A**) and GVAFTWVMA (residues 156-164) at the N-terminal end of TM4 (**Fig. 8B**), exhibited clear EX1-like behavior in ligand-free opsin and in the presence of all three non-retinoid ligands (results for quercetin only are shown). This indicates a local cooperative opening and closing dynamics at these regions. Although deuterium uptake was quantified at only two time points (0 and 10 min), precluding full kinetic modeling, multiple lines of evidence strongly support the presence of genuine EX1-like exchange in these regions. These include: (i) a pronounced redistribution of the isotopic envelope after 10 min, characterized by a marked reduction in low-mass (less deuterated) species and a concomitant increase in a highly deuterated population ^36,40^; (ii) robust bimodality of the isotope clusters, with F-test P values < 0.01 for binomial fits, indicating that a single Gaussian distribution cannot adequately describe the spectra (**Fig. 8**, **plots (2) and (3)**); and (iii) confirmation of bimodal behavior by resampling analysis ^41^, reducing the likelihood that the observed bimodality arises from noise or undersampling rather than true kinetic heterogeneity. Classic HDX-MS studies on proteins such as cytochrome c and pepsin have used similar patterns, bimodal envelopes, loss of low-mass species, and time-dependent population shifts, to identify EX1-like behavior associated with cooperative unfolding transitions ^36,40^.

**Figure 8.**
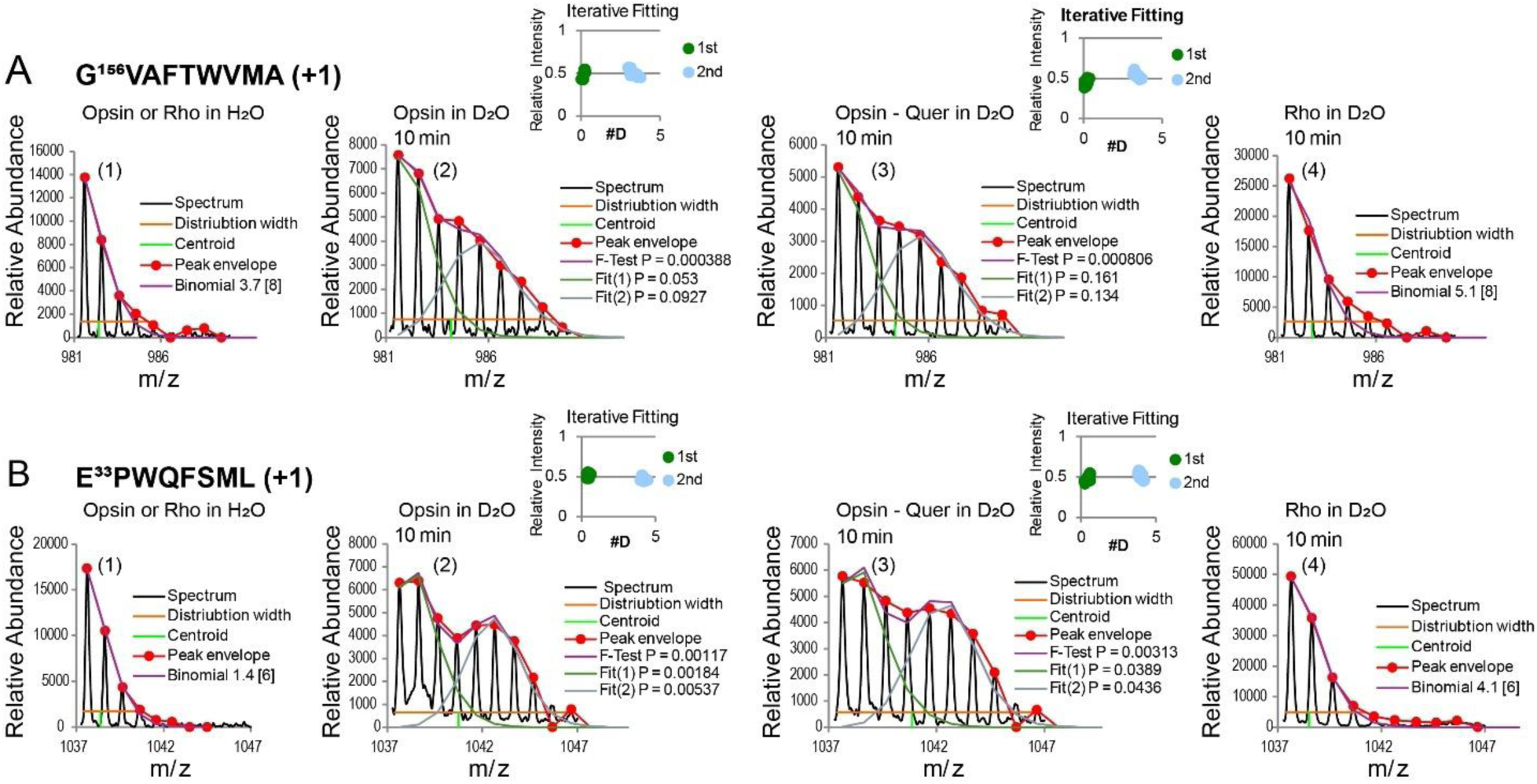
Binomial fitting of deuterated mass envelops for N-terminal segments of TM1 and TM4. **A**, Amino acid sequence and corresponding mass envelope for TM4 N-terminal peptide. **B**, Amino acid sequence and corresponding mass envelope for TM1 N-terminal peptide. Single binomial fits are shown in the leftmost (1) and rightmost (4) panels for opsin/rhodopsin (Rho) control sample in H_2_O and Rho in D_2_O, respectively. The number of amides used for the best fit (violet) is indicated in the square brackets. The P_F-test_ values (comparing unimodal and bimodal fits) are indicated in the figure. The plots numbered (2) and (3) show the double binomial fit for ligand-free opsin and opsin-quercetin complex in D_2_O, respectively. The corresponding P-value of the intensity parameters for each population is shown in green and gray. The small inset panels in plot (2) and (3) show the results from resampling analysis, where each point represents a deuteration level (x axis) and relative intensity (y axis).

Notably, this EX1-like feature was completely suppressed upon binding of 11-*cis*-retinal (**Fig. 8, plot (4)**), consistent with a shift toward a more uniformly packed ensemble, whereas non-retinoid compounds failed to eliminate this behavior (**Fig. 8, plot (3)**). Although in the presence of quercetin, the lower N-terminal segment of TM1 exhibits decreased deuterium uptake, this localized stabilization is not strong enough to eliminate the EX1-like behavior, likely because it fails to propagate across TM1 as seen upon binding of 11-*cis*-retinal (**Fig. 2A**). To mechanistically interpret EX1-like behavior, we employed protein PSN analysis, which revealed that ligand binding induces rewiring of intramolecular interaction networks and the formation of distinct residue interaction hubs that correlate with changes in conformational dynamics (**Fig. 9A**). In the case of 11-*cis*-retinal binding new hubs are formed, centered at Tyr^29^, Tyr^43^, Met^44^, and Met^183^ near N-terminus of TM1, as well as centered at Phe^148^, Trp^161^, Cys^167^, Tyr^206^, and His^211^ near N-terminus of TM4. However, based on our previous MD simulation-derived PSN analysis ^15^, quercetin binding induces the formation of only two new interaction hubs localized to the N-terminal regions of TM1 (centered at Tyr^29^) and TM4 (centered at Phe^148^), which are shared with retinal-induced hubs, and one distinct hub in N-terminal region of TM2 (centered at Pro^71^), while leaving the remainder of this local network largely unchanged (**Fig. 9B**). Therefore, this reorganization of the intramolecular interaction network likely explains how retinal binding stabilizes the otherwise flexible N-terminal regions of TM1 and TM4 and suppresses EX1-like exchange behavior, an effect that non-retinal compounds fail to achieve.

**Figure 9.**
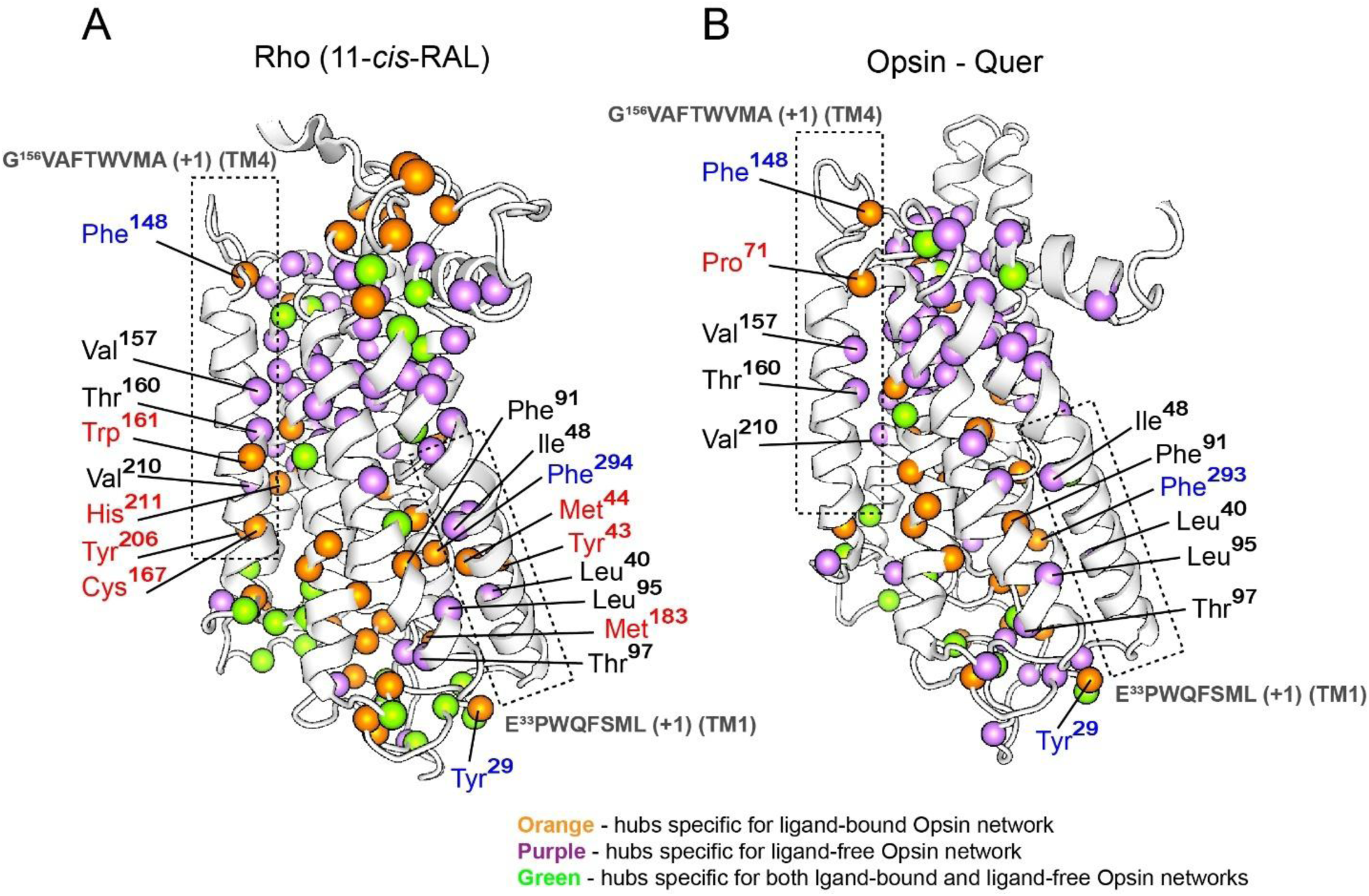
Protein Structure Network (PSN) for peptides with EX1-like kinetics. **A**, Difference PSNs between ligand-free opsin (PDB ID: 3CAP) and rhodopsin (Rho) (PDB ID: 1U19). **B**, Difference PSNs between ligand-free opsin (PDB ID: 3CAP) and quercetin-bound opsin ^15^. The N-terminal peptides of TM4 (left) and TM1 (right) are shown in dotted boxes. Hubs specific to liganded opsin and unliganded opsin networks are colored orange and purple, respectively, while those shared by both networks are colored green. This data highlights the differences in intramolecular communication hubs after binding each ligand. Residues forming ligand-induced hubs shared by 11-*cis*-retinal and quercetin are shown in blue, while residues forming distinct ligand-specific hubs are shown in red.

## DISCUSSION

While the biological effects and binding properties of non-retinoid pharmacological chaperones have been extensively studied through biochemical and cell-based assays, there is limited direct, high-resolution structural evidence from techniques like X-ray crystallography or cryo-EM for their interaction with rod opsin. In this study, using amide-HDX and His-HDX-MS alternative structural approaches, we have provided empirical structural details for the interactions of rod opsin with non-retinoid pharmacochaperons, such as flavonoids (quercetin and myricetin), and CR5 chromenone. Here, we compare the percent deuterium uptake by rod opsin upon binding of its natural ligand 11-*cis*-retinal to form rhodopsin, with that observed upon binding of non-retinoid pharmacochaperones, flavonoids or CR5, to both unliganded opsin and rhodopsin. By analyzing these differences, we provide first structural insight into ligand-specific interactions within the rod visual receptor.

### Long-Range Structural Rearrangement upon Orthosteric Binding of Endogenous Ligand

Our HDX data revealed that binding of 11-*cis*-retinal to ligand-free opsin induces a pronounced decrease in deuterium uptake across TM4-TM7 and multiple loop segments, indicating a global reduction in solvent accessibility and structural fluctuations (**Fig. 2**). This behavior is consistent with the long-standing view of 11-*cis*-retinal as an endogenous inverse agonist that constrains rhodopsin in a low-entropy, inactive conformation and suppresses basal activity ^42–45^. Crystallographic and spectroscopic studies established that chromophore, covalently linked to Lys^296^ as a protonated Schiff base, engages an extensive hydrogen-bond and packing network that spans TM3, TM5-TM7 and couples to conserved E(D)RY and NPxxY motifs on the cytoplasmic side ^4,46,47^. Our HDX data complement these static structures by directly demonstrating how introducing 11-*cis*-retinal into flexible opsin drives the ensemble toward a more conformationally restricted state, with protection in TM4-TM7, arguing that retinal binding does not simply fill a local cavity but reshapes the helical packing landscape.

Decreased deuterium uptake in the N-terminal region, ICL1, ECL1, and portions of ICL2, ICL3, and ECL2 show that this stabilization propagates outward from the orthosteric site, through the disulfide-linked ECL2 and the extracellular helical ring, into the intracellular loops that define the G-protein recognition surface. Such long-range propagation has been widely hypothesized based on GPCR structures and simulations ^48^ but has rarely been mapped experimentally at this resolution. Recent HDX-MS work on the β₁-adrenergic receptor has shown that ligand-induced conformational fingerprints can distinguish agonists, antagonists and biased ligands, and can be used to prioritize early hits and guide structure activity relationship optimization in GPCR drug discovery ^49^. The convergence of the HDX changes with alterations in residue interaction network and hubs identified by our PSN analysis provides an important mechanistic bridge (**Figs. 3, 9, and Supplementary Fig. 1**). PSN approaches treat residues as nodes and persistent non-covalent contacts as edges, allowing the identification of highly connected hubs and shortest interaction networks that underline intramolecular signaling ^28^. Previous works applying PSN to rhodopsin and related GPCRs, showed that activation, dimerization and misfolding rewire hub patterns and path efficiencies between extracellular and intracellular poles ^15,17,48,50^. Our data shows that segments with increased HDX protection upon 11-*cis*-retinal binding, notably TM4-TM7 and specific loop regions, substantially overlap with residues whose hub scores and participation in critical communication pathways shift between the opsin and rhodopsin PSN models (**Fig. 3**). This implies that retinal binding stabilizes the nodes and edges that carry intramolecular information, restoring a canonical inactive-state communication backbone rather than merely rigidifying a local pocket.

The combination of backbone amide-HDX and His-specific HDX further links global stabilization to local side-chain microenvironments. His^65^, His^100^, and His^152^ become more protected in rhodopsin, whereas His^195^ shows a decreased His-HDX half-life (**Fig. 2**), implicating ECL2 and its Zn-binding site in state-dependent tuning of the intradiscal surface ^26,51–53^. Notably, His^195^ could not be assigned in the amide-HDX map, highlighting the complementarity of the two probes. Totally, consistent trends across backbone and His-HDX support a model in which retinal binding reorganizes an extended hydrogen-bonding and hydration network that couples helices, loops, and key hubs. This distributed stabilization provides a structural basis for the exceptional efficacy of 11-*cis*-retinal and 9-*cis*-retinal as pharmacological chaperones for diverse rhodopsin mutants, at least *in vitro* ^12,17,54,55^. Because receptor stability and signaling depend on the integrity of this interconnected network, disease-causing mutations or pharmacological ligands that perturb these structural hubs can exert disproportionate effects, even when they are distant from the retinal-binding pocket or the G protein-coupling interface. This interpretation is consistent with recent PSN-based analyses of multiple adRP mutants, which mapped misfolding propensity onto network hubs and identified regions amenable to small-molecule rescue ^17,56^.

### Structural Clues for Orthosteric Binding of Flavonoids and CR5

Our amide and His specific HDX data (**Fig. 2A**) indicate that quercetin, myricetin, and chromenone CR5 all engage the same orthosteric cavity as 11-*cis*-retinal in ligand-free opsin, but with more locally confined structural consequences. Deuterium uptake decreases in helices surrounding the retinal pocket, and the pattern of protection closely parallels with that seen upon 11-*cis*-retinal binding. This is in agreement with docking analysis performed in this study (**Table 1**, **Fig. 6, and Supplementary Figs. S2 and S3**) and previous studies ^13,16^ showing that these non-retinoid compounds can occupy the retinal-binding pocket of rod opsin. However, in contrast to 11-*cis*-retinal which form covalent protonated Schiff base with Lys^296 57^, quercetin, myricetin and CR5 bind non-covalently and lack the same rigid, shape-complementary fit, as shown in previous docking ^13,16^. This explains the lower magnitude of HDX protection for flavonoids and CR5, relative to 11-*cis*-retinal. Notably, myricetin and CR5 additionally increased deuterium uptake in TM1 and more distal regions in contrast to the uniformly stabilizing effect of 11-*cis*-retinal. This suggests that some non-retinoids introduce local structural perturbations or redistribute flexibility away from the deep pocket toward the periphery of the helical bundle. Such local softening could reflect alternative hydrogen-bond networks or changes in lipid/protein contacts, especially given that TM1 participates in inter-helical networks linking TM1-TM2-TM7 via residues such as Leu^40^, Thr^58^, Leu^76^, and Phe^91 57^ as observed in the PSN analysis (**Fig. 3**).

The His-HDX data provides a second, more fine-grained layer of evidence that these non-retinoid ligands occupy the orthosteric site but engage the surrounding network differently. In rhodopsin, His^65^ and His^152^ lie near the cytoplasmic region, while His^100^ and His^195^ are present in intradiscal motifs that can bind Zn²⁺ ^20,51,57^. His^65^, His^152^ and His^211^ also participate in the Meta-I/Meta-II equilibrium. As previously reported, mutation of His^65^ or His^152^ enhances proton sensitivity of the Meta-I/Meta-II transition, whereas His^211^ strongly stabilizes Meta-II upon protonation ^58^. Quercetin-bound opsin showed the most retinal-like His-HDX signature (**Fig. 2B**). This suggests that quercetin recapitulates the retinal-induced microenvironmental shift at the cytoplasmic (His^65^ and His^152^) sites but only weakly reorganizes the intradiscal surface, consistent with docking that places quercetin in the deep pocket with possible extension toward a TM5-TM6-ECL2 site ^13,21^. Functionally, this aligns with quercetin’s behavior as a potent stabilizer and pharmacological chaperone that improves opsin folding and P23H mutant homeostasis without perfectly duplicating dark-state rhodopsin ^13,15^. In contrast, myricetin and CR5 produce inverted His fingerprints relative to retinal, with only partial overlap (**Fig. 4**). This implies that myricetin stabilizes the cytoplasmic region in a manner broadly similar to 11-*cis*-retinal, while reorganizes the intradiscal region in a distinct way, potentially rendering it more solvent-exposed or dynamically labile. However, CR5 drives a different electrostatic and hydration pattern toward both the cytoplasmic side (His^65^) and the intradiscal Zn²⁺-coordinating region (His^100^/His^195^). Together, His-HDX provides a microenvironment barcode for each ligand explaining their structural rational behind their different pharmacochaproning properties ^13,16,21^. Quercetin exhibits most retinal-like properties, myricetin promotes a more opened intradiscal architecture, and CR5 selectively stabilizes the core pocket while rebalancing distal surfaces.

### Structural Clues for Allosteric Modulatory Effects of Flavonoids and CR5

Our amide-HDX and His-HDX data, together with acid denaturation assay and molecular docking to cytoplasmic cavity, also reveals genuine allosteric modulatory behavior of non-retinoid ligands for the dark state conformation. In 11-*cis*-retinal-bound rhodopsin, myricetin and CR5 induce a broad increase in deuterium uptake across multiple TMs, with localized protection near residues 85-90 and the cytoplasmic end of TM4 (**Fig. 5A**). This pattern is consistent with a shift toward a more conformationally dynamic ensemble rather than simple global unfolding, suggesting formation or stabilization of specific contacts near TM4. In contrast, quercetin induces more modest structural rearrangements. Accordingly, myricetin and CR5 behave as stronger cytoplasmic allosteric modulators that partially loosen the intracellular interface, whereas quercetin exerts a weaker, more localized effect, analogous to how G proteins, arrestins, and nanobodies reshape the GPCR intracellular cavity to tune active and inactive substates ^1,54,59–61^. This differential behavior may help explain the previously observed increase in rhodopsin regeneration rates in the presence of quercetin ^21^ and other flavonoids ^62^.

His-HDX in dark-adapted rhodopsin supports this view at the side-chain level. On its own, 11-*cis*-retinal stabilizes the dark state and produces a characteristic pattern of His-HDX *t*_1/2_ ^20,26^ that reflects a specific electrostatic and hydration environment around His^65^, His^100^, His^152^, and His^195^ (**Fig. 5B**). Quercetin decreases *t*_1/2_ at His^65^, His^100^, and His^152^, indicating a generally softened microenvironment at both cytoplasmic and extracellular/intradiscal sites, consistent with the modest HDX changes observed. Myricetin increases *t*_1/2_ at His^65^ but decreases it at other His residues, suggesting that it stabilizes the cytoplasmic side near His^65^ in agreement to amide-HDX. CR5 increases *t*_1/2_ at His^195^ but decreases it at other His residues, indicating preferential stabilization of the ECL2/intradiscal region and a loosening at cytoplasmic and central transmembrane regions, which is in contrast to amide-HDX result. These ligands specific His fingerprints indicate that each compound, while binding in a similar cytoplasmic region, couples differently into the global electrostatic and hydration network of rhodopsin. This level of detail goes beyond earlier work that inferred allosteric modulation largely from functional readouts ^21^. This study now maps these allosteric outcomes onto specific His residues and regions, providing a structural rationale for their distinct pharmacological profiles. These structural changes correlate with functional readout, as all three ligands accelerated acid-induced decay of rhodopsin (**Fig. 4**) in the order myricetin > CR5 > quercetin, consistent with their allosteric K_d_ values and the magnitude of HDX perturbations. Docking these ligands into the cytoplasmic cavity of the dark-state rhodopsin (PDB ID: 1U19) further supports experimental observations. Quercetin and myricetin preferentially occupy the central region of this cavity formed between TM2, TM3, TM4, TM6, and TM7, whereas CR5 adopts multiple poses that span this cavity and extent into adjacent interhelical grooves (**Fig. 7**), consistent with its broader HDX footprint.

Overall, these observations are fully consistent with earlier reports showing that flavonoids can modulate rhodopsin activation and regeneration yet typically exert only modest effects on downstream signaling under physiological conditions ^13–15,21^. The dual behavior we observe, chaperoning apo-opsin while subtly loosening the dark-state conformation of retinal-bound rhodopsin, is well aligned with established principles of GPCR allosteric pharmacology, in which the functional impact of a ligand depends strongly on the receptor’s initial conformational and ligand-bound state ^61,63–65^. Importantly, this context dependence suggests a mechanistic basis for selectively stabilizing misfolded or apo states without substantially perturbing native photopigment function. Accordingly, our findings provide design principles for next-generation allosteric modulators and pharmacological correctors that could target mechanistically distinct classes of rhodopsin mutations or enhance mutant selectivity while minimizing interference with pigment regeneration at the orthosteric pocket.

### 11-cis-retinal Suppresses EX1-like Behavior by Stabilizing Local Gates and Global Packing

Our amide-HDX data indicated that the flexible ends of TM1 (residues 33-40) and TM4 (residues 156-164) exhibit EX1-like signatures in ligand-free opsin (**Fig. 8**) suggests that these segments function as local gating elements that occasionally undergo relatively large-amplitude opening events, likely coupled to partial rearrangements of the helical bundle. A striking feature of our data is that the EX1-like hallmark completely disappears upon 11-*cis*-retinal binding. This indicates that the conformational fluctuation regime of these segments is fundamentally changed in the rhodopsin state. Mechanistically, this can be concluded that upon binding of 11-*cis*-retinal, the local packing geometry around the N-terminal ends of TM1 and TM4 and their connections to adjacent helices are tightened, such that the higher-entropy open state is no longer significantly populated on the HDX timescale. The exchange regime thus reverts to EX2-like behavior, with slow, incremental uptake and no distinct ‘all-or-none’ exchange events. This interpretation aligns with our broader finding that 11-*cis*-retinal induces widespread protection across TM4-TM7 and reconfigures PSN communication hubs (**Figs. 3 and 9**). In particular, PSN analysis reveals that new and unique hubs (**Fig. 9A**) arise in rhodopsin and lie adjacent to the EX1-reporting peptides (E^33^PWQFSML and G^156^VAFTWVMA), which are enriched in hydrophobic and aromatic residues. These new hubs are located at Tyr^29^, Met^44^, Tyr^43^, Met^183^, and Phe^294^ residues around TM1, and Phe^148^, Trp^161^, His^211^, Tyr^206^, and Cys^167^ residues around TM4 segments. We propose that these hubs represent new stabilizing contact networks that lock the ends of TM1 and TM4 into more ordered states, selectively quenching rare cooperative opening pathways at strategic positions.

Thus, the stabilization of local gates (at TM1/TM4 termini in opsin) and global packing (across TM4-TM7) give a reference to interpret whether new ligands mimic the endogenous stabilization pattern or create a different one. In contrast, all the non-retinoid ligands in this study exhibited a partial improvement in global packing. In addition, these non-retinoid ligands failed to remove the cooperative opening signature at specified local gates (at TM1/TM4 termini in opsin), as indicated by backbone HDX data (**Fig. 8, plot (3)**), likely due to distinct reorganizations of the intramolecular interaction network. As shown in **Fig. 9B**, although quercetin binding induces new communication hubs around TM1 (Tyr^29^ and Phe^293^ residues) and TM4 (Pro^71^ and Phe^148^ residues), it does not fully recapitulate the hub architecture established by 11-*cis*-retinal. This could possibly explain why these ligands, unlike retinal, stabilize a more limited range of disease related mutations ^16,54^. Together, these observations show that chromophore binding does more than stabilize the pocket, it actively reshapes the fluctuation landscape of specific structural elements. This establishes global packing and local gate stabilization as a benchmark ensemble for effective pharmacochaperoning. From a drug-design perspective, ligands that suppress EX1-like behavior emerge as explicit targets for more effective pharmacochaperones.

## CONCLUSION

High-resolution structural determination of GPCR-ligand complexes remains technically challenging. Therefore, we employed complementary hydrogen-deuterium exchange (HDX) strategies to gain structural insight into rhodopsin-non-retinoid ligand interactions. The combined use of segment-resolved amide-HDX and residue-specific His-HDX provides a powerful approach to define both ligand binding sites (orthosteric versus peripheral) and the propagation of structural and electrostatic changes throughout the receptor. Using this platform, we show that flavonoids and CR5 do more than simply occupy the orthosteric site of rod opsin: their binding actively reshapes the conformational fluctuation landscape of specific structural elements. In the presence of 11-*cis*-retinal, these ligands additionally engage a cytoplasmic allosteric pocket, remodel structural dynamics and His microenvironments, and accelerate acid-induced pigment decay in a ligand-specific manner. Furthermore, the long-range structural rearrangements induced by 11-*cis*-retinal provide a mechanistic explanation for how opsin is converted into a rigid, inactive rhodopsin pigment, why the native chromophore and its analogues are exceptionally effective pharmacological chaperones, and which structural interaction hubs must be stabilized to maintain GPCR integrity. The discovery of EX1-like behavior at the N-terminal ends of TM1 and TM4, and its complete suppression by 11-*cis*-retinal, reveals that the natural chromophore not only globally stabilizes the receptor but also eliminates cooperative opening pathways at key helical termini.

Together, these findings define core principles of non-retinoid pharmacochaperoning, clarify retinal’s dual role as an inverse agonist and pharmacological chaperone, and establish a general framework for linking mutation-specific destabilization, ligand-induced structural rescue, and functional outcomes. These concepts are likely to extend beyond rhodopsin to other class A GPCRs, where small molecules modulate receptor folding and signaling by reshaping conformational ensembles.

## EXPERIMENTAL PROCEDURES

### Chemical Reagents

n- Dodecyl-β-D-maltoside (DDM) was purchased from Affymetrix Inc. (Maumee, OH). Dimethyl sulfoxide (DMSO), 9-*cis*-retinal were obtained from Sigma (St. Louis, MO). EDTA-free protease inhibitor cocktail tablets were purchased from Roche (Basel, Switzerland). Immobilizes pepsin was purchased from Thermo Scientific (USA). Pepsin powder was purchased from Worthington Biovhemical Corp. (Lakewood, NJ, USA). Deuterated water (D_2_O) was obtained from Cambridge Isotop Laboratories, Inc (Andover, MA, USA). Myricetin (PubChem CID 5281672) and quercetin (PubChem CID 5280343) were purchased from Sigma. Chromenone CR5 (PubChem CID 42220126) was purchased from OTAVAchemicals, Ltd (Ontario, Canada).

### Isolation of Rod Outer Segments and Preparation of Opsin Membranes

Bovine rod outer segment (ROS) membranes from frozen retinas were isolated under dim red light using sucrose gradient centrifugation as described previously ^66^. The isolated ROS membranes were washed with a buffer composed of 5 mM HEPES pH 7.5, 1 mM EDTA, and 1 mM DTT followed by centrifugation at 25,000xg for 25 min. This procedure was repeated four times. Final membrane pellet containing rhodopsin was stored at -80 °C for future experiments or was resuspended in the buffer composed of 10 mM sodium phosphate pH 7.5, and 20 mM hydroxylamine, and exposed to white light with a 150-Watt bulb for 1 h at 0 °C to prepare opsin membranes. Membranes were then pelleted by centrifugation at 16,000xg for 10 min, followed by wash with 10 mM sodium phosphate, pH 6.5 and 2% BSA four times. Subsequently these membranes were washed with 10 mM sodium phosphate pH 6.5 four times. Final membrane pellet was resuspended in 20 mM bis-tris propane (BTP), pH 7.5 and 120 mM NaCl and stored at -80 °C for future experiments.

A UV-Vis spectrophotometer (Cary 60, Varian, Palo Alto, CA) and the absorption coefficients ε_500_ nm=40,600 M^−1^cm^−1^ and ε_280_ nm=81,200 M^−1^cm^−1^ were used respectively to measure the concentration of rhodopsin and opsin ^67,68^.

### Purification of Opsin and Rhodopsin using Affinity Chromatography

ROS membranes or opsin membranes were resuspended in 20 mM BTP pH 7.5 containing 120 mM NaCl and 20 mM DDM and incubated for 1 h at 4 °C on a nutator to solubilize the membranes, followed by centrifugation at 16,000xg for 1 h at 4 °C. The soluble fraction was combined with 250 µl of 2 mg 1D4/ml agarose beads (anti-Rho C-terminal 1D4 antibody immobilized on cyanogen bromide (CNBr)-activated agarose) and incubated for 1 h at 4 °C on a nutator. Next, the beads were transferred to a column and washed with 10 ml of 20 mM BTP, pH 7.5, 120 mM NaCl, containing 2 mM DDM. Rhodopsin or opsin were eluted with the same buffer, supplemented with 0.6 mg/ml of the 1D4 peptide (TETSQVAPA) and 0.4 mM DDM, aliquoted and stored at -80 °C for future experiments. All the purification steps were carried out in dark conditions. The protein concentrations were measured with the UV-Vis spectrophotometer as described above. The samples were concentrated by Amicon filter 30K, to reach a concentration ∼1.5 mg/ml, then aliquoted in 20 µl final volume and stored at -80 °C before amide-HDX experiment.

### Amide-HDX of Rhodopsin, Opsin and Their Ligand-Complexes

Amide-HDX exchange experiments were performed and analyzed using previously published protocol with modifications ^20^. Samples (∼30 µg) of purified opsin or rhodopsin, or their complexes with quercetin, myricetin or chromenone CR5 were diluted in 80 µl of D_2_O and kept at room temperature for 10 min to achieve steady state exchange. To obtain receptor ligand complexes 1 µl of compound was added to the receptor samples with concentration ratio 1:1 (∼40 µM each) and incubated for at least 60 min before HDX exchange was initiated. To terminate the reaction, the ice-cold quenching buffer was added, composed of 50% formic acid either in H_2_O or D_2_O (to deuterate samples), to reach pH 2.5, followed by addition of pepsin (Worthington, Lakewood, NJ, USA) at 6 mg/ml (5 µl). The samples were digested on ice for 10 min. Next, the samples were loaded on a C18 20 x 2 mm Luna column (Phenomenx, Torrance, CA, USA) with a temperature-controlled Hewlett-Packard autosampler set to 4 °C. Peptides were eluted with the following gradient sequence: 0-4 min, 98% H_2_O with 0.1% (v/v) formic acid (A) and 2% acetonitrile with 0.1% formic acid (B); from 0-30 min, 98% to 2% A. Separation was achieved at flow rate 0.3 ml/min using Agilent 1100 HPLC system (Agilent Technologies, Santa Clara, CA USA). The eluent was directed into an LTQ Velos linear Trap mass spectrometer (Thermo Scientific, Watman, MA, USA) via electrospray ionization source operated in a positive ion mode. Peptides were identified by tandem mass spectrometry (MS^2^), employing the Xcalibur version 4.0.27.19 (Thermo Scientfic) to extract the relative signal intensity as a function of m/z and comparing with theoretical MS-Digest. The data was deconvoluted with HXExpress-3v and the average deuterium content of each fragment ion was calculated by using the centroid of its isotopic cluster. HDX exchange was color-coded based on the maximal observed deuterium uptake and represented as percentage of total theoretical uptake or change in the percentage of deuterium uptake between opsin and rhodopsin, and their ligand flavonoid/chromenone complexes.

### Histidine-HDX of Rhodopsin, Opsin and Their Ligand-Complexes

Opsin and rhodopsin in outer segment membranes were used. Briefly, membranes were resuspended in 20 mM BTP, pH7.5, 120 mM NaCl to a final concentration of 1 mg/ml. Ligands were added into the sample at protein: ligand ration 1:50 and incubated for at least 1 h on ice followed by centrifugation at 16,000xg for 30 min to pellet the membranes. Then, pellet was resuspended in D_2_O buffer composed of 20 mM BTP, pH 7.5, 120 mM NaCl to a final volume of 600 µl, aliquoted into three 200 µl samples, incubated at 37 °C for 70 h. To stop D_2_O exchange reactions, each 200 µl samples were centrifuged at 16,000xg for 1 min, the supernatant was discarded, while membrane pellet was solubilized in 20 µl of 1 % SDS in 0.1 % aqueous HCOOH. Five volumes of cold (-20 °C) acetone were added, and samples were stored for 1 h at -20 °C to precipitate protein. The precipitates were centrifuged and washed three times in cold acetone (90%) followed by incubation in a bath type sonicator used to re-suspend the protein pellet. These protein pellets were solubilized in 50 µl of 1:5 CH3OH:HCOOH mixed with 10 µl of fresh performic acid (5% v/v H_2_O_2_ in HCOOH). Incubation of these mixtures on ice for 30 min allowed to oxidize the cysteine and methionine to cysteic acid and methionine sulfone residues, respectively, a step which enables recovery of peptides containing each of the His residues in rhodopsin. Oxidized protein was dried using Speedvac and resuspended in 10-30 µl of neat HCOOH, diluted with H_2_O to reach 6% HCOOH, aliquoted into 50 µl and stored at -20 °C.

Each 50 µl sample was digested with gentle agitation for 60 min at 25 °C with 5 µl of immobilized pepsin (Princeton Separations). All above steps were performed at dark conditions for rhodopsin. The digested peptides desalted using BioPureSPN mini C18 cartridge, and finally dissolved in 30 µl of 0.1% formic acid before sending for LC-MS/MS.

Peptides were analyzed using LC-MS/MS UltiMate 3000 LC system (Dionex, San Francisco, CA) interfaced to a LTQ-Orbitrap XL mass spectrometer (Thermo-Finnigan, Breman, Germany). The platform was operated in the nano-LC mode using the standard nano-ESI API stack fitted with a picotip emitter (uncoated fitting, 10 µm spray orifice, New Objective Inc., Woburn, MA). Separation was performed at a flow rate of 300 nl/min with a 1:1,000 splitter system. Five µl of protein digests were injected into a C18 PepMap trapping column (0.3 x 5 mm, 5 µm particle size, Dionex) equilibrated with 0.1% trifluoroacetic acid (TFA) and 2% CH3CN (v/v) and washed for 5 min with the equilibration solvent at a flow rate of 25 µl/min, by using an isocratic loading pump operated through an autosampler. After washing, the trapping column was switched in-line with a C18 Acclaim PepMap 100 column (0.075 x 150 mm, Dionex) and peptides were separated with a linear gradient of 2% to 50% CH3CN in aqueous 0.1% HCOOH over a period of 40 min at 300 nl/min with direct introduction of the eluate into the mass spectrometer. The mass spectrometer was operated in a data-dependent MS to MS/MS switching mode, with the five most intense ions in each MS scan subjected to MS/MS analysis. The full MS scan was performed at 60,000 resolution and the subsequent MS/MS analysis was performed at 30,000 resolution. The total scan cycle frequency was about 1 sec. The precursor ion isolation width was set at m/z ± 2.0, allowing transmission of the M and M^+2^ isotopic ions of the peptide for Collision-induced dissociation (CID). The threshold intensity for the MS/MS trigger was set at 2,000 and fragmentation was carried out in the CID mode with a normalized collision energy (NCE) of 35. Data were collected in profile mode. The dynamic exclusion function for previously selected precursor ions was not enabled during these analyses. Xcalibur version 4.0.27.19 (Thermo Scientifics) was used for instrument control, data acquisition, and data processing.

### Acid denaturation of dark-adapted rhodopsin at presence of flavonoids

Rhodopsin in outer segment membranes at a final concentration of 2.5 µM were incubated in buffer composed of 20 mM BTP, pH 7.5, 120 mM NaCl, and 6 mM DDM for 30 min at 4 °C on a nutator under dark conditions. The sample centrifuged at 16,000xg for 15 min, to separate any undissolved precipitation. Clear supernatant then incubated with each ligand at a serial dilution 1/3: 0, 0.062, 0.19, 0.56, 1.67, 5, and 15 µM for 30 min. Then 50 µl HCl 300 mM were added to each sample to reach a final HCl concentration of 30 mM and stable pH ∼ 2-3, and immediately the UV-Vis spectrum of samples was acquired at 1 min time interval. The change in absorption intensity at λ_500_ nm were recorded. The normalized native conformation fraction, X(t), computed using the equation (I), and each trace fitted to equation (II), to calculate the reaction rate, K_obs_.

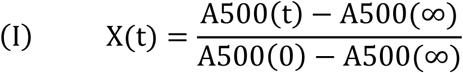

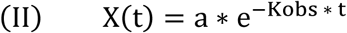

In which, X(t) is native conformation fraction at time t. A_500_(0), A_500_(t), A_500_(∞) are absorption intensity at time zero (i.e. when acid is not added), at time t and after the end of reaction, respectively. To calculate the K_d_ for allosteric interactions of each ligand, calculated K_obs_ at each ligand dilution vs ligand concentration fitted to the equation (III):

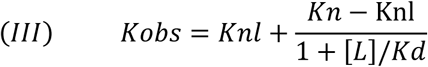

In which, [L] is ligand concentration. K_n_ and K_nl_ are the reaction rate with no ligand ([L]=0) and at saturating ligand concentration (here [L]=15 µM) respectively.

### Docking Simulation and Bioinformatics Evaluation

Docking simulations of ligands with rod opsin models were performed using the Aoutodock vina to identify important amino acid residues for the ligand-receptor interaction. For orthosteric position, local docking performed using PDB IDs: 3CAP and 2I36 to model apo-receptor respectively in active and inactive state. For the allosteric position, local docking was performed using PDB ID: 1U19, which contains a complete C-terminal region at inactive state. One monomeric unit of the protein was selected, and the co-crystallized molecules and all crystallographic water molecules were removed from the coordinate set; hydrogen atoms were added, and partial charges were assigned to all atoms. The 3D structure of the compounds used in docking were downloaded from PubChem or generated using Avogadro and their geometry were optimized. Before docking the ligand-free structure of opsin model was subjected to a 10 ns molecular dynamics equilibration with Memprot. GPCR-ModSim web server (https://memprot.gpcr-modsim.org/). Then, the molecular docking of ligands into the opsin orthosteric binding pocket or cytoplasmic region was performed.

Protein Structure Network analysis (PSN) performed using webPSN webserver ^27^. We applied PSNs difference mode using PDB ID: 3CAP and 1U19 respectively as ligand-free opsin and inactive rhodopsin conformational state, to investigate the Links, Hubs, and Global MetaPath.

### Statistical Analysis

Data was obtained from at least three independent experiments. Statistical significance was assessed using Student’s *t*-test, with P < 0.05 considered significant. Error bars represent the standard deviation (S.D).

## AUTHORSHIP CONTRIBUTIONS

*Participated in research design:* B.J., Z.P., J.T.O., M.M., and M.G.

*Conducted experiments:* B.J., Z.P., J.T.O., M.M., and M.G.

*Performed data analysis:* B.J., Z.P., M.M., and M.G.

*Wrote or contributed to the writing of the manuscript:* B.J., Z.P, J.T.O., M.M., and M.G.

## ACKNOWLEDGEMENTS

This research was supported by the National Institutes of Health (NIH) (R01 EY032874 to B.J., R01 EY023948 to M.G.) and by the NIH P30 grant (P30EY011373, the Visual Sciences Research Center Core Facilities at Case Western Reserve University), and by the NIH Shared Instrument S10 Grant S10OD023436-01 for the Thermo Scientific Fusion Lumos LC-MS/MS. We thank Dr. Belinda Willard, Director of the Cleveland Clinic Lerner Research Institute’s Proteomics and Metabolomics Core for performing mass spectrometry analysis of His-HDX samples.

## DATA AVAILABILITY STATEMENT

The data that support the findings of this study are included in the manuscript.

