## Supplemental Information for "Conformational signatures of native ligand and pharmacochaperone binding in rhodopsin"

**RUNNING TITLE:** Non-retinoid Modulators of Rhodopsin

[*] **CORRESPONDENCE**

Beata Jastrzebska, Ph.D., Department of Pharmacology, School of Medicine, Case Western Reserve University, 10900 Euclid Ave., Cleveland, OH 44106-4965, USA; Phone: 216-368-5683; Fax: 216-368-1300;, ORCID ID: https://orcid.org/0000-0001-5209-8685.

**Keywords:** GPCR Rhodopsin; Pharmacochaperones, Hydrogen-Deuterium Exchange Mass Spectrometry: HDX-MS; Allosteric Modulation; Non-retinoid Ligands

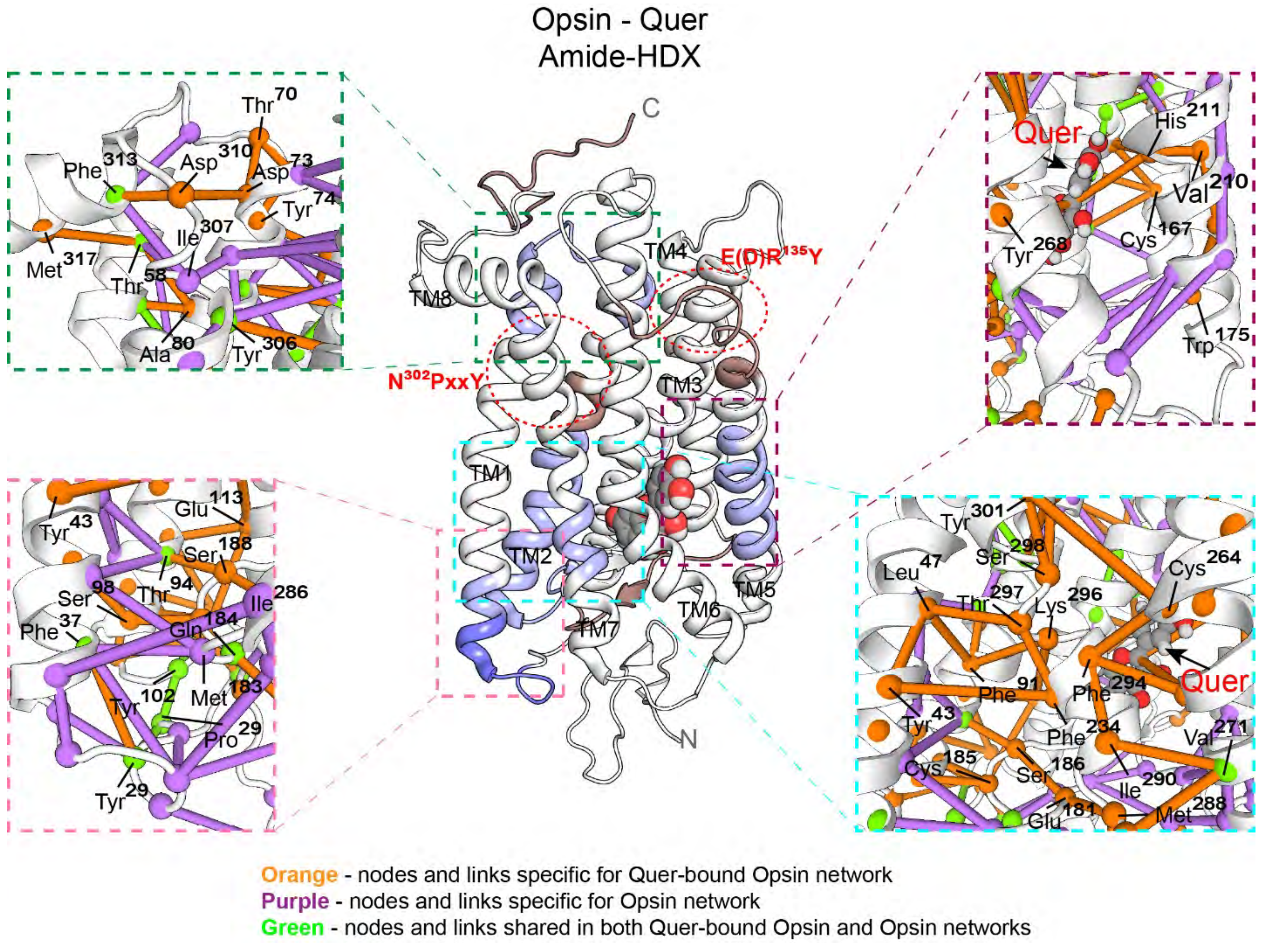

**Supplementary Figure 1. Structural amide-HDX pattern for quercetin-bound rhodopsin and corresponding Protein Structure Network (PSN) models.** The structural rearrangement model for opsin-quercetin amide-HDX is shown in middle panel. The intensity of blue color displays the percentage of deuterium uptake reduction (decreased solvent accessibility). Locations of NPxxY, and E/DRY motifs are indicated by red dashed ovals. Quercein is shown with white-red-brown colored spheres. The Global MetaPath derived from PSN analysis corresponds to each structural segments are shown in separate boxes. Nodes and links specific to ligand-free opsin (PDB ID: 3CAP) and quercetin-bound opsin (15) networks are represented by purple and orange color respectively. Nodes and links shared by both networks are colored green. Generation of new MetaPaths after quercetin binding (orange color) are in good concordance with change in percentage of deuterium uptake across transmembrane region, suggesting corresponding network and structural rearrangement following endogenous ligand binding.

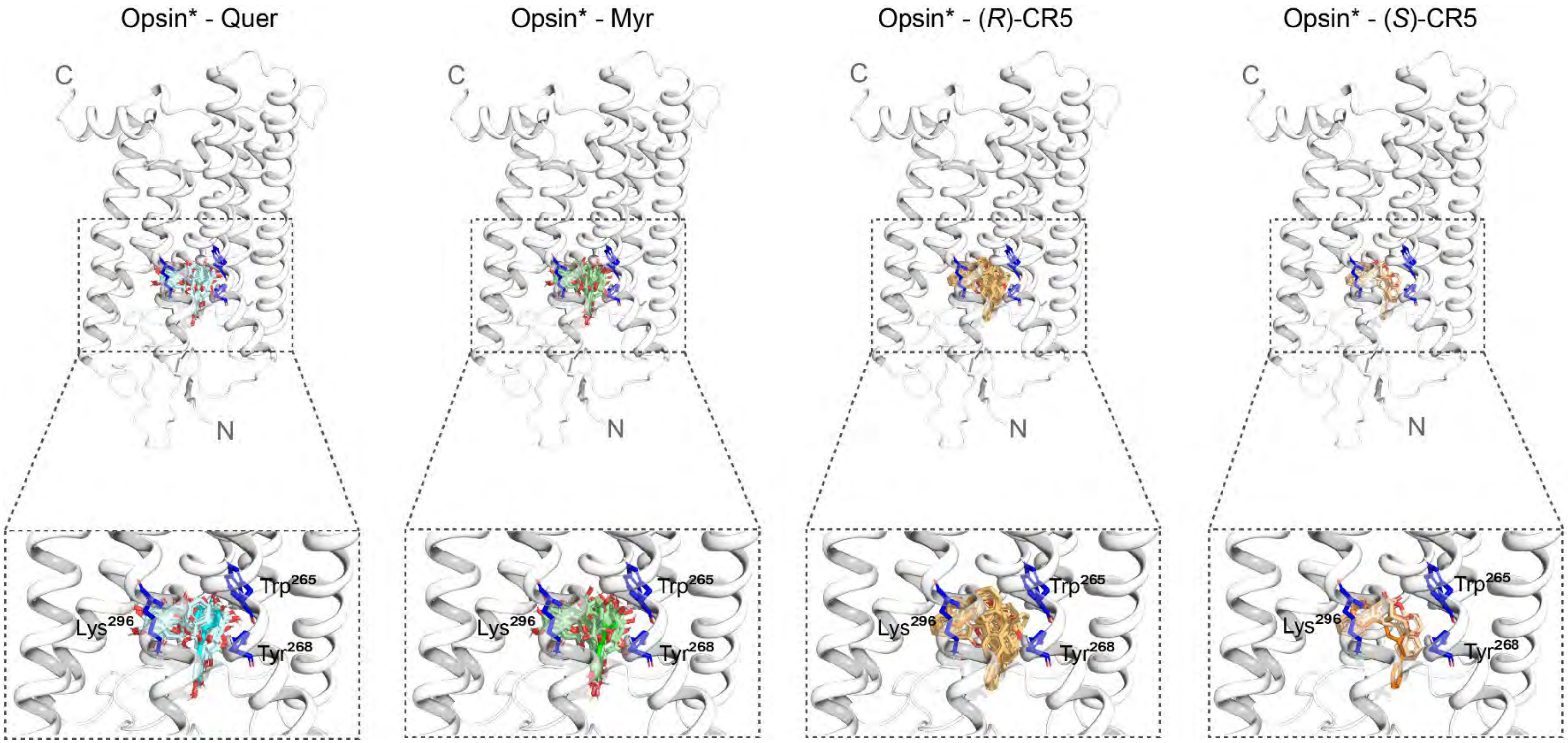

**Supplementary Figure 2. Local interactions between bovine rod opsin and the flavonoids or CR5 within the orthosteric pocket.** Molecular docking analysis of ligand interactions within the orthosteric pocket of rod opsin using structure of apo-opsin with active conformation (Opsin*) (PDB ID: 3CAP). Docking was performed using AutoDock Vina in PyRx environment. The top panel shows all identified docking poses for each ligand ranked by docking score. And the bottom panel shows zoomed-in views for all the docking poses within the orhosteric site with the best-ranked poses highlighted with brighter colors.

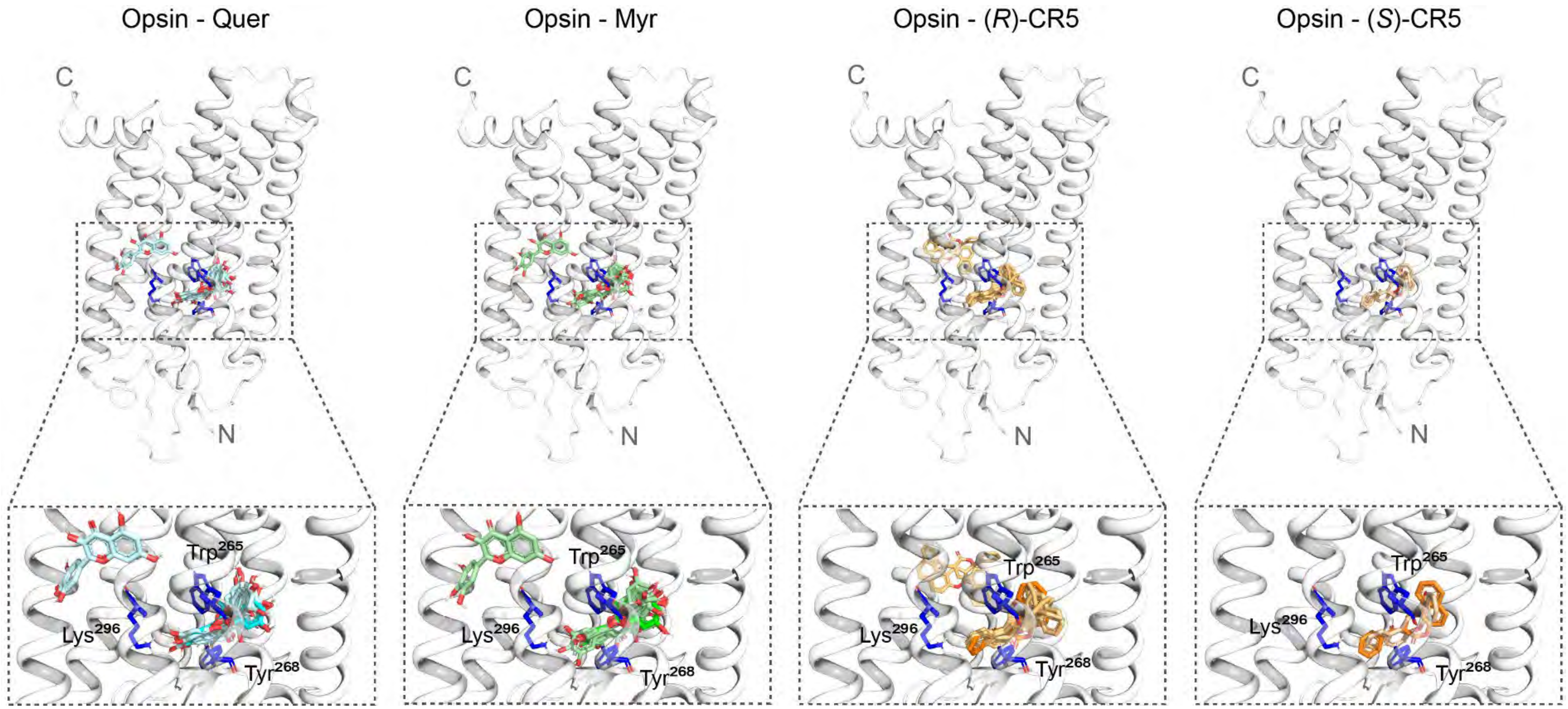

**Supplementary Figure 3. Local interactions between bovine rod opsin and the flavonoids or CR5 within the orthosteric pocket.** Molecular docking analysis of ligand interactions within the orthosteric pocket of rod opsin using structures of apo-opsin with inactive conformation (Opsin) (PDB ID: 2I36). Docking was performed using AutoDock Vina in PyRx environment. The top row shows all identified docking poses for each ligand ranked by docking score. And the bottom row shows zoomed-in views for all the docking poses within the orhosteric site with the best-ranked poses highlighted with brighter colors.

**Supplementary Table 1. Sequences of peptic fragments of rod opsin showing normalized deuterium uptake for rod opsin and rhodopsin with different ligands bound**

| | | | | | Δ %Deuterium Uptake | | | | | | |
|---|---|---|---|---|---|---|---|---|---|---|---|
| | | | | | 11-*cis*-RAL ortosteric effect | Q ortosteric effect | Q allosteric effect | M ortosteric effect | M allosteric effect | CR5 ortosteric effect | CR5 allosteric effect |
| Peptide Sequence | Charge | m/z | Max | Real Max | %D Rho %D Ops | %D Ops/Q - %D Ops | %D Rho/Q - %D Rho | %D Ops/M - %D Ops | %D Rho/M - %D Rho | %D Ops/CR5 - %D Ops | %D Rho/CR5 - %D Rho |
| V[11]PFSNKTGVVRS PFEAPQYY | 2+ | 1691.27 | 16 | 12.8 | -8.00 | -1.44 | -0.32 | -2.50 | 2.50 | -2.04 | 1.01 |
| Y[29]LAEPWQF | 1+ | 1053.504 | 6 | 4.8 | -10.01 | -5.57 | -1.34 | 1.82 | 6.17 | -0.38 | 5.32 |
| Y[29]LAEPWQFSML | 2+ | 692.830 | 9 | 7.2 | -8.94 | -0.32 | 1.42 | 3.40 | 3.43 | -0.53 | 0.59 |
| Y[29]LAEPWQFSML | 1+ | 1384.661 | 9 | 7.2 | -10.48 | -0.09 | 1.08 | 0.86 | 2.55 | -1.06 | 0.47 |
| L[47]IMLGFPINF | 1+ | 1164.649 | 8 | 6.4 | -12.45 | -0.26 | -0.65 | 2.64 | 1.33 | -2.78 | -1.16 |
| L[45]LIMLGFPINF | 1+ | 1277.733 | 9 | 7.2 | -10.91 | 0.22 | -2.49 | 5.44 | 8.38 | 8.44 | 19.44 |
| M[49]LGFPINFLTL | 2+ | 633.348 | 9 | 7.2 | -9.77 | 0.02 | -0.40 | 4.71 | 3.97 | 4.00 | 4.05 |
| Y[60]VTVQHKKLRTPL NYIL | 2+ | 1043.609 | 15 | 12 | -6.70 | -2.50 | -1.23 | -2.62 | 0.71 | -4.56 | -0.36 |
| I[75]LLNL | 1+ | 585.397 | 4 | 3.2 | -11.89 | -0.17 | 0.39 | 1.69 | 2.09 | 0.96 | 1.49 |
| Y[74]ILLNL | 1+ | 748.574 | 5 | 4 | -13.73 | 0.63 | 0.47 | 2.96 | 0.86 | 5.06 | 1.17 |
| L[77]NLAVADLF | 1+ | 975.551 | 8 | 6.4 | -19.67 | -2.12 | 2.15 | 2.45 | 5.47 | -0.27 | 1.24 |
| A[80]VADLF | 1+ | 635.370 | 5 | 4 | -10.09 | 0.56 | 0.09 | 4.70 | 3.02 | 6.03 | 4.53 |
| F[85]MVFGGF | 1+ | 804.374 | 6 | 4.8 | -4.60 | 0.25 | -0.53 | -1.50 | -1.28 | -1.49 | -1.36 |
| M[86]VFGGFTTTL | 2+ | 537.266 | 9 | 7.2 | -5.36 | -1.52 | -1.12 | -0.97 | -0.37 | -1.50 | 0.52 |
| T[93]TLYTSLHGYFVF | 1+ | 1031.760 | 12 | 9.6 | -6.07 | -3.77 | 0.23 | 10.97 | 19.91 | 8.11 | 17.71 |
| H[100]GYFVF | 1+ | 769.366 | 5 | 4 | -11.32 | -1.38 | -2.01 | -3.06 | 8.93 | -5.82 | 12.14 |
| A[117]TLGGE | 1+ | 547.400 | 5 | 4 | -3.88 | 0.26 | -0.22 | 0.21 | 4.12 | 1.98 | 5.77 |
| T[118]LGGEIALWSLV VLAIERY | 2+ | 1052.090 | 18 | 14.4 | -2.19 | -0.01 | 0.17 | -0.15 | 1.69 | -0.21 | 1.61 |
| A[124]LWSLVVL | 1+ | 900.555 | 7 | 5.6 | -3.30 | 0.90 | 1.00 | 2.11 | 3.38 | 2.72 | 2.61 |

| | | | | | | | | | | | |
|---|---|---|---|---|---|---|---|---|---|---|---|
| R147FGENHAIMGVA | 2+ | 651.321 | 11 | 8.8 | -5.36 | -1.42 | 2.00 | -5.13 | -5.78 | -4.06 | -6.04 |
| F159TWVMALACAA | 1+ | 1183.564 | 10 | 8 | -4.28 | -2.39 | -0.02 | 0.38 | 3.95 | 1.60 | 7.39 |
| L165ACAAPPLVGW | 1+ | 1097.581 | 8 | 6.4 | -11.43 | 1.86 | -0.34 | -14.27 | -0.59 | -0.52 | 0.52 |
| A169PPLVGWSRYI | 1+ | 1258.694 | 8 | 6.4 | -7.26 | 1.10 | -0.77 | -0.49 | 3.11 | -0.29 | 4.65 |
| W175SRYIPEGMQ | 1+ | 1266.690 | 8 | 6.4 | -9.55 | 2.01 | -1.29 | 4.45 | 3.68 | 5.75 | 1.70 |
| V204IYMF | 1+ | 672.342 | 4 | 3.2 | -12.34 | -0.56 | -1.67 | 4.76 | 4.27 | 3.34 | 9.35 |
| M207FVVHF | 1+ | 779.390 | 5 | 4 | -20.25 | -1.76 | 0.87 | 10.72 | 5.52 | 12.21 | 4.24 |
| V209VHFII | 1+ | 727.450 | 5 | 4 | -14.94 | 0.12 | -0.60 | 1.98 | 1.39 | -0.10 | 1.40 |
| Y206MFVVHFII | 1+ | 1168.622 | 8 | 6.4 | -19.73 | -3.49 | -0.58 | -5.73 | 1.59 | -2.19 | -0.20 |
| Y206MFVVHFIIPL | 1+ | 1378.75 | 9 | 7.2 | -12.96 | -2.79 | 0.44 | -2.79 | 0.44 | -3.50 | -0.79 |
| I217VIF | 1+ | 491.322 | 3 | 2.4 | -17.18 | 0.74 | 3.01 | -3.62 | 11.71 | -4.53 | 12.42 |
| F221CYGQLVF | 2+ | 488.729 | 7 | 5.6 | -2.69 | -0.99 | -2.22 | 2.90 | 2.38 | 0.41 | 1.80 |
| C222YGQLVFTVKEA | 1+ | 1357.682 | 11 | 8.8 | -8.55 | 2.51 | 0.15 | -1.24 | 1.96 | -1.12 | 1.35 |
| E247KEVTRMVII | 1+ | 1217.692 | 9 | 7.2 | -7.64 | 0.64 | -0.06 | 12.75 | 20.53 | 8.68 | 6.43 |
| E247KEVTRMVIIMVI AFL | 2+ | 946.537 | 15 | 12 | -5.89 | -0.27 | -1.28 | -0.42 | 2.75 | -1.57 | 2.11 |
| F261LICWLPY | 1+ | 1054.543 | 6 | 4.8 | -13.30 | 1.55 | -0.03 | -3.27 | 2.84 | -0.24 | 2.58 |
| F287MTIPA | 1+ | 679.348 | 4 | 3.2 | -10.26 | 1.68 | 0.33 | -1.17 | 0.64 | -1.07 | 0.81 |
| P291AFFA | 1+ | 552.280 | 3 | 2.4 | -15.44 | -1.34 | 0.31 | 15.60 | 17.77 | 27.85 | 18.73 |
| A295KTSAVYNPVIY | 2+ | 663.355 | 10 | 8 | -9.98 | 0.62 | 1.31 | -5.19 | -4.65 | -5.82 | -4.59 |
| F294AKTSAVYNPVIY | 1+ | 1472.778 | 11 | 8.8 | -8.74 | 0.23 | 1.12 | 2.30 | 7.78 | 2.39 | 7.25 |
| P291AFFAKTSAVYN PVIY | 2+ | 894.468 | 13 | 10.4 | -11.72 | -0.05 | 0.10 | -3.07 | 8.08 | 1.61 | 10.55 |
| I307MMNKQF | 1+ | 911.450 | 6 | 4.8 | -20.04 | 1.06 | -0.14 | 0.09 | 2.64 | -0.17 | 1.85 |
| M308MNKQF | 1+ | 798.360 | 5 | 4 | -17.57 | 1.86 | -0.42 | 2.51 | 3.30 | 6.81 | 4.96 |
| M308MNKQFRNCMV TTL | 2+ | 858.902 | 13 | 10.4 | -10.51 | 1.24 | 1.89 | -1.12 | 5.61 | -1.11 | 4.52 |
| G329DDEA | 1+ | 506.172 | 4 | 3.2 | -6.64 | 0.18 | 0.67 | 6.19 | 8.74 | 10.07 | 9.59 |
| S334TTVSKTETSQV APA | 2+ | 753.882 | 13 | 10.4 | 1.93 | 3.47 | -1.13 | 7.53 | 3.09 | 5.38 | 0.25 |

Column 1 shows peptic peptides sequence from the rod opsin primary sequence. Column 2 indicates the charge of the ion. Column 3 exhibits the *m/z* of the ion used to identify the peptide based on the MS/MS spectrum. Column 4 shows the maximum number of theoretically exchangeable sites of the peptide fragment (Max = number of non-proline peptide bonds in the peptide fragment). Column 5 reveals the uptake normalized to 80% of the theoretical maximum exchangeable sites. The 80% normalization reflects the dilution percentage in $D_2O$. Columns 6-12 shows the difference of deuterium uptake percentage between each two specified states. The numbers colored in green indicate statistically significant changes, $P < 0.05$ calculated with the Student's *t*-tests. Green: $P < 0.05$; Orange: $0.08 > P > 0.05$; Black: $P > 0.08$. Ops, opsin; Rho, rhodopsin; Q, quercetin; M, myricetin.

**Supplementary Table 2. Half-lives of His-HDX for membrane-bound rod opsin and rhodopsin-flavonoids or CR5 complexes**

| | half-lives (h) of His-HDX for different histidine | | | | | | | | | | | |
|---|---|---|---|---|---|---|---|---|---|---|---|---|
| | **H65 (YVTVQHKKLRTPLN *m/z* 566.3)** | | | | **H100 (HGYFVFGPTGCNLEGFF *m/z* 970.4)** | | | | **H152 (HAIMGVAF *m/z* 439.2)** | | | |
| | **OM** | S.D. | **ROS** | S.D. | **OM** | S.D. | **ROS** | S.D. | **OM** | S.D. | **ROS** | S.D. |
| **$D_2O$** | **12.98** | 0.73 | **14.6** | 0.46 | **20.57** | 0.62 | **33.49** | 10.18 | **62.84** | 2.95 | **154.92** | 53.79 |
| **Quercetin in $D_2O$** | **13.54** | 0.46 | **13.26** | 0.51 | **20.62** | 0.43 | **24.56** | 0.47 | **116.61** | 18.54 | **93.67** | 16.89 |
| **Myricetin in $D_2O$** | **13.73** | 0.73 | **17.49** | 2.28 | **16.44** | 1.29 | **25.28** | 4.85 | **44.51** | 1.19 | **73.71** | 2.35 |
| **CR5 in $D_2O$** | **11.79** | 0.07 | **12.21** | 0.04 | **19.78** | 0.82 | **18.04** | 0.51 | **95** | 6.77 | **82.21** | 3.23 |
| | **H195 (YYTPHEETNNESF *m/z* 815.84)** | | | | **H211 (VVHFIIPL *m/z* 469.3)** | | | | **H278 (YIFTHQGSDFGPIF *m/z* 814.8)** | | | |
| | **OM** | S.D. | **ROS** | S.D. | **OM** | S.D. | **ROS** | S.D. | **OM** | S.D. | **ROS** | S.D |
| **$D_2O$** | **7.55** | 0.56 | **7.2** | 0.002 | **35.42** | 7.64 | **71.91** | 80.58 | **103.21** | 34.69 | **232.84** | 137.95 |
| **Quercetin in $D_2O$** | **6.7** | 0.48 | **7.22** | 0.47 | **61.75** | 18.25 | **43.31** | 6.28 | **380.05** | 347.53 | **152.06** | 23.95 |
| **Myricetin in $D_2O$** | **9.05** | 0.45 | **7.23** | 0.25 | **23.69** | 2.01 | **36.01** | 12.29 | **147.34** | 30.06 | **949.11** | 498.58 |
| **CR5 in $D_2O$** | **7.91** | 0.12 | **8.39** | 0.29 | **49.75** | 21.48 | **52.63** | 34.64 | **-226.14** | 397.45 | **73.77** | 4.81 |

The peptide sequences and corresponding *m/z* values used to calculate histidine exchange half-lives are shown. Half-lives were calculated from three independent replicates, and statistical significance was assessed using Students *t*-tests. For $H^{211}$ and $H^{278}$, the standard deviation (S.D.) was high due to substantial variability among replicates. Therefore, these residues were excluded from the final evaluation. ROS, Rhodopsin in Membrane; OM, Opsin in Membrane; S.D., Standard Deviation.

### Supplementary Table 3. Ligand Docking to Rod Opsin or Rhodopsin

| | | Active Opsin PDB ID: 3CAP | | | | Inactive Opsin PDB ID: 2I36 | | | | Rhodopsin PDB ID: 1U19 | | | |
|---|---|---|---|---|---|---|---|---|---|---|---|---|---|
| | | **Q** | **M** | **(R)-CR5** | **(S)-CR5** | **Q** | **M** | **(R)-CR5** | **(S)-CR5** | **Q** | **M** | **(R)-CR5** | **(S)-CR5** |
| **Type of interactions** | **Dock Scores for the best pose** | -9.6 | -9.6 | -9.7 | -10.4 | -7.8 | -7.8 | -9.3 | -11.3 | -7 | -7.3 | -8.2 | -8 |
| | **Hydrophobic** | Glu181 (3.41 Å)<br>Tyr192 (3.74 Å)<br>Trp265 (3.96 Å) | Thr118 (3.95 Å)<br>Glu181 (3.38 Å)<br>Tyr192 (3.77 Å) | Ala117 (3.52 Å)<br>Thr118 (3.62 Å)<br>Glu181 (3.80 Å)<br>Tyr268 (3.95 Å)<br>Ala292 (3.97 Å)<br>Ala296 (3.93 Å)<br>Ala296 (3.90) | Ala117 (3.46 Å)<br>Glu181 (3.29 Å)<br>Tyr192 (3.61 Å)<br>Ala292 (3.80 Å) | Phe212 (3.79 Å)<br>Trp265 (2.70 Å)<br>Tyr268 (3.73 Å)<br>Tyr268 (3.30 Å) | Phe212 (3.79 Å)<br>Trp265 (2.70 Å)<br>Tyr268 (3.73 Å)<br>Tyr268 (3.26 Å) | Ala117 (3.81 Å)<br>Thr118 (3.97 Å)<br>Phe261 (3.42 Å)<br>Trp265 (3.07 Å)<br>Trp265 (3.59 Å)<br>Trp265 (3.09 Å)<br>Tyr268 (3.80 Å)<br>Ala292 (3.30 Å)<br>Ala295 (3.79 Å) | Ala117 (3.76 Å)<br>Phe261 (3.24 Å)<br>Trp265 (3.28 Å)<br>Trp265 (3.87 Å)<br>Tyr268 (3.25 Å)<br>Tyr268 (3.84 Å)<br>Ala292 (3.32 Å) | Ala246 (3.56 Å)<br>Lys311 (3.46 Å) | Thr70 (3.74 Å) | Thr70 (3.76 Å)<br>Leu72 (3.77 Å)<br>Val250 (3.40 Å)<br>Lys311 (3.58 Å)<br>Gln312 (3.96 Å) | Tyr74 (3.86 Å)<br>Tyr74 (3.71 Å)<br>Ile154 (3.82 Å)<br>Trp161 (3.94 Å)<br>Trp161 (3.58 Å) |
| | **Hydrogen Bonds** | Thr118 (2.62 Å)<br>Gly121 (3.35 Å)<br>Glu181 (3.42 Å)<br>Gly188 (3.68 Å)<br>Tyr192 (2.02 Å)<br>Tyr192 (2.00 Å)<br>Pro291 (2.76 Å) | Glu181 (3.43 Å)<br>Gly188 (3.72 Å)<br>Tyr192 (1.98 Å)<br>Tyr192 (1.99 Å)<br>Pro291 (2.79 Å) | Ile189 (2.89 Å) | Ile189 (2.19 Å) | Thr118 (3.64 Å)<br>Thr118 (2.26 Å)<br>Thr118 (2.28 Å)<br>Gly121 (3.42 Å)<br>Glu122 (2.32 Å)<br>Glu122 (3.50 Å)<br>Cys187 (1.68 Å)<br>His211 (2.04 Å)<br>Lys296 (3.29 Å) | Thr118 (3.62 Å)<br>Thr118 (2.25 Å)<br>Thr118 (2.28 Å)<br>Gly121 (3.43 Å)<br>Glu122 (2.32 Å)<br>Glu122 (3.51 Å)<br>Cys187 (1.68 Å)<br>His211 (2.03 Å)<br>Ala292 (3.15 Å)<br>Lys296 (3.29 Å) | Ile189 (2.18 Å)<br>Tyr268 (1.89 Å) | Ile189 (2.37 Å)<br>Tyr191 (2.73 Å)<br>Tyr268 (2.61 Å) | Ala234 (3.54 Å)<br>Gln237 (2.65 Å)<br>Lys311 (2.04 Å)<br>Lys311 (2.5 Å)<br>Gln312 (2.13 Å)<br>Gln312 (2.09 Å) | Glu134 (2.13 Å)<br>Glu249 (2.26 Å)<br>Glu249 (2.10 Å)<br>Glu249 (3.04 Å)<br>Asn310 (2.30 Å)<br>Asn310 (2.16 Å)<br>Gln312 (3.28 Å) | Gln312 (2.54 Å)<br>Ser334 (2.01 Å) | // |

| | | | | | | | | | | | | | |
|---|---|---|---|---|---|---|---|---|---|---|---|---|---|
| | **π-Stacking** | Tyr191 (4.77 Å) Tyr268 (5.49 Å) | Tyr191 (4.74 Å) | Tyr191 (4.95 Å) | Tyr191 (4.81 Å) | // | // | Tyr268 (3.97 Å) | // | // | // | // | Trp161 (3.97 Å) Trp161 (3.89 Å) Trp161 (3.68 Å) |
| | **π-Cation Interactions** | Lys296 (4.79 Å) | Lys296 (4.88 Å) | Lys296 (4.93 Å) | // | // | // | // | // | // | // | // | // |

The details of the interactions between each non-retinoid ligands and rod opsin side chains within the orthosteric binding pockets (PDBs 3CAP and 2I36) or cytoplasmic cavity in rhodopsin (PDB 1U19). Q, quercetin, M, myricetin.
